# Quantifying and correcting bias in transcriptional parameter inference from single-cell data

**DOI:** 10.1101/2023.06.19.545536

**Authors:** Ramon Grima, Pierre-Marie Esmenjaud

## Abstract

The snapshot distribution of mRNA counts per cell can be measured using single molecule FISH or single-cell RNA sequencing. These distributions are often fit to the steady-state distribution of the two-state telegraph model to estimate the three transcriptional parameters for a gene of interest: mRNA synthesis rate, the switching on rate (the on state being the active transcriptional state) and the switching off rate. This model assumes no extrinsic noise, i.e. parameters do not vary between cells, and thus estimated parameters are to be understood as approximating the average values in a population. The accuracy of this approximation is currently unclear. Here we develop a theory that explains the size and sign of estimation bias when inferring parameters from single-cell data using the standard telegraph model. We find specific bias signatures depending on the source of extrinsic noise (which parameter is most variable across cells) and the mode of transcriptional activity. If gene expression is not bursty then the population averages of all three parameters are overestimated if extrinsic noise is in the synthesis rate; underestimation occurs if extrinsic noise is in the switching on rate; both underestimation and overestimation can occur if extrinsic noise is in the switching off rate. We find that some estimated parameters tend to infinity as the size of extrinsic noise approaches a critical threshold. In contrast when gene expression is bursty, we find that in all cases, the mean burst size (ratio of the synthesis rate to the switching off rate) is overestimated while the mean burst frequency (the switching on rate) is underestimated. We estimate the size of extrinsic noise from the covariance matrix of sequencing data and use this together with our theory to correct published estimates of transcriptional parameters for mammalian genes.

## I. INTRODUCTION

The two-state model of gene expression [1, 2], also called the telegraph model, provides a simple framework describing the stochastic nature of gene expression. The model consists of four effective reactions:

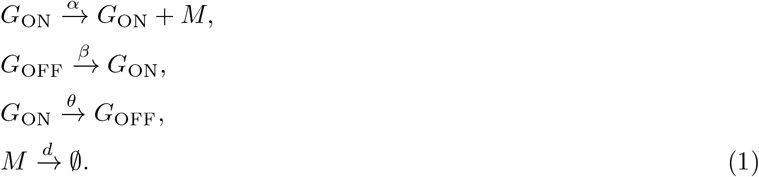

These reactions describe mRNA (M) synthesis in the active promoter state G_ON_ with rate α, switching from the inactive promoter state G_OFF_ to the active state with rate β, switching from the active to the inactive promoter state with rate θ and mRNA degradation with rate d respectively. Assuming Markovian dynamics, the telegraph model can be solved in steady-state leading to an analytical marginal distribution of mRNA counts [1].

It has been shown that the transcript count distributions for many genes, obtained from a snapshot population measurement using smFISH (single molecule fluorescence in situ hybridization) or scRNA-seq (single cell RNA sequencing), can be well fit by the steady-state distributions predicted by the telegraph model [3–5]. This matching also simultaneously leads to estimates of the three main transcriptional parameters normalised by the degradation rate, i.e. α/d, β/d and θ/d. Note that without an explicit measurement of the mRNA degradation rate d, it is not possible to obtain the absolute values of the other parameters. Estimation of these parameters provides insight into the mode of transcriptional activity and its relationship with core promoter elements (such as gene length, TATA and initiator elements)[6]. For example a switching on rate that is much larger than a switching off rate would indicate that expression occurs almost incessantly, whereas the reverse condition together with a large synthesis rate suggests expression occurs in large infrequent bursts.

Despite its simplicity and utility, there are doubts about the telegraph model’s ability to capture the major biochemical processes that influence the observed stochasticity in mRNA numbers [7–12]. Gene expression is known to be affected by various processes that are not explicitly modelled by the telegraph model, e.g. cell cycle dynamics [4, 13], coordination of transcription with cell size [14, 15], mitochondrial dynamics [16, 17] and Ca^2+^ signalling [18, 19]. These various processes (and others) can be grouped together under the umbrella of extrinsic noise, since they arise independently of a gene of interest but act on it. Some studies have concluded that for many mammalian genes, the variability of cytoplasmic transcript abundance is determined primarily by the phenotypic cellular state [19, 20]. Thus correcting data for extrinsic noise before it is input to an inference framework based on the telegraph model is important since potentially this type of noise could significantly bias parameter estimation.

In principle, the dual reporter method enables discrimination between extrinsic noise and intrinsic noise (locally determined by the gene sequence and which is modelled by the telegraph model) [21]. This has been done for some organisms using fluorescent protein reporters [22–24] but the experiments are technically challenging and they do not directly enable the study of transcription since mRNA and protein are not generally strongly correlated [25]. There are also parameter inference methods that instead of fitting to population data, they fit to individual cells using single-cell time-lapse protein data [26, 27]. However live-cell imaging approaches which allow visualization of mRNA bursts as they occur in living cells are challenging since they are low-throughput and require genome-editing [28, 29]. An alternative approach to correct the parameter estimates for noise sources not taken into account by the telegraph model is to directly eliminate these sources from the data. For example, to correct for the fact that the telegraph model assumes one gene copy, whereas in reality some cells will have double the gene copy number than others, we can separate the data according to cell-cycle phase. Then we can fit a telegraph model to single allele data from cells in G1 and separately fit the distribution obtained from a convolution of the distributions of two telegraph models to double allele data from cells in G2 [13] (under the simple assumption that copies are independent of each other). This procedure has the added benefit of accounting for cell-cycle dependence of transcriptional parameters [4, 30, 31] which is not described by the telegraph model. While it is still not widespread, this type of correction can nowadays be readily performed since cell-cycle phase classification is possible for both smFISH data (most of the experiments already include a DNA-labelled channel) [13] and scRNA-seq data (deep learning applied to the unspliced-spliced RNA space of cell cycle-related genes) [32].

However these corrections do not go far enough because there are additional noise sources. For example, different cells within the same cell-cycle phase may have different effective synthesis rates of cytoplasmic mRNA because of the coupling of gene expression to cell size for some eukaryotic genes [14, 19] and cell-to-cell variation in the nuclear export rate [33]. While some of these sources could be accounted for by carefully designed experiments to measure extrinsic noise sources of choice (such as cell volume) the fact is that presently we are far from having a complete understanding of what are the most relevant sources. Hence it stands to reason that after any correction we may do to partition the data to exclude one noise source or another, inevitably there will still be cell-to-cell variability in the transcriptional parameters due to unaccounted noise sources [34–38] and fitting the telegraph model to this data may lead to biased estimates. A summary of the standard fitting procedure is illustrated in Fig. 1. *A main question of interest that remains unresolved is: what type (positive / negative) and extent of bias in parameter estimation should we expect when fitting the telegraph model to experimental data?* In the best case scenario, one hopes that the inferred parameters are to a good approximation equal to their true values, i.e. the parameter values averaged over all observed cells.

**FIG. 1.**
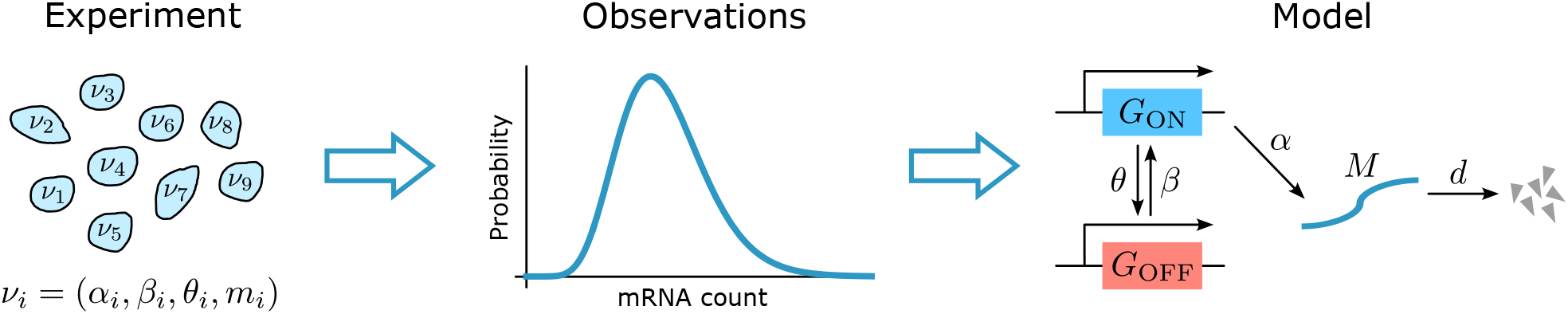
Illustration of the standard transcriptional parameter inference procedure. In a population of cells, the transcriptional rates for a particular gene vary from cell to cell: *α*_*i*_, *β*_*i*_ and *θ*_*i*_ are the mRNA synthesis rate, the switching on rate and the switching off rate for cell *i*, respectively. It is assumed that the degradation rate does not vary. The mRNA count in cell *i* is denoted by *m*_*i*_. In a typical inference setup, the steady-state distribution of mRNA (*M*) per cell is measured and then by fitting this to the analytical steady-state distribution of the standard telegraph model, the three transcriptional rates (normalised by the degradation rate) for the gene are estimated. Since the telegraph model assumes the parameters are the same for each cell, it is unclear what is the relationship of the inferred parameters to the true population averaged parameters.

To address this question, here we develop a moment-based analytical approach to understand how the size and sign of estimation bias depends on the mode of transcriptional activity, the source of extrinsic noise (which parameter varies most from cell-to-cell) and the type of distribution that describes it. We identify specific “bias signatures” which we confirm by inferring parameters, using the method of maximum likelihood, from simulated single cell data. In particular, we show that while the telegraph model distributions can well fit those computed from data, the inferred parameters from the telegraph model can be far away from their true population average. We also use the theory to correct published telegraph model estimates of the mammalian transcriptome using scRNA-seq data [6].

## II. THE SOURCE OF EXTRINSIC NOISE AND TIMESCALE SEPARATION DETERMINE THE SIZE AND SIGN OF ESTIMATION BIAS

Generally all of the parameters of the telegraph model may vary from cell to cell and their spread is described by some arbitrary joint distribution. However because of its generality this is difficult to study and hence we make some simplifying assumptions. We consider three different cases: (i) variability only in the synthesis rate ρ; (ii) variability only in the switching on rate k_on_; (iii) variability only in the switching off rate k_off_. Note that we have given the rate parameters different labels than in the reaction scheme (1) to distinguish them from those in the standard telegraph model where there is no parameter variability between cells. Note that we have not considered variability in the mRNA degradation from one cell to another, because this is difficult to study – since the steady-state telegraph model solution is a function of the three transcriptional rates normalised by the degradation rate, variability in the degradation rate would imply variability in all three inferred rates simultaneously. In all cases the distribution of each parameter (normalised by the degradation rate) is assumed to be a gamma distribution – this distribution is ideal because it is defined on the positive real line and its mean and variance can be tuned independently. The three circuits are illustrated on the left hand side of Fig. 2. Our question is: if for a gene of interest the distribution of mRNA counts per cell for each of these three cases is fitted to that of the standard telegraph model, how are the parameters inferred related to the true population averaged parameters? For example if the mRNA distribution comes from a telegraph model where the varying parameter is the synthesis rate then it would be of interest to understand what is the relationship of the population averaged synthesis rate to the synthesis rate inferred from fitting the data to the standard telegraph model.

**FIG. 2.**
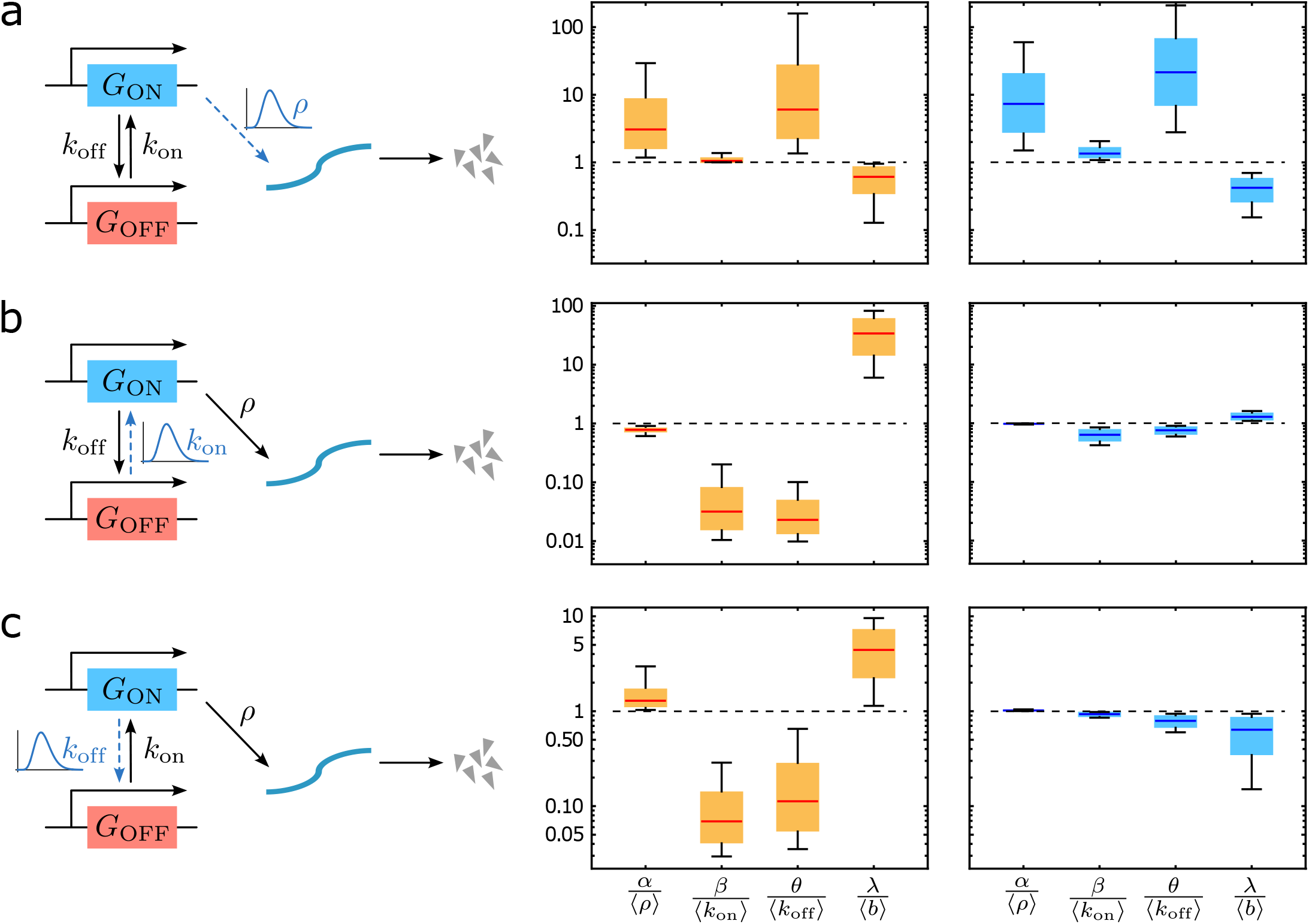
Systematic bias in parameter estimation due to unaccounted extrinsic noise sources. We consider cell populations with (a) variable synthesis rates, (b) variable switching on rates and (c) variable switching off rates. The variability in a parameter is modelled by a gamma distribution. For each case, we generate a million random parameter sets and estimate the effective parameters of the standard telegraph model (reaction scheme (1)) by exactly matching the first three moments of this model to those of telegraph model with extrinsic noise. For those parameters sets where all inferred parameters are positive, the statistics of their values normalised by the ground truth values (the population averaged parameters) are shown by box-whisker plots in the middle column of the figure (lower and upper fences show the 10th and 90th percentiles, respectively; the yellow bar shows the interquartile range; median is shown by the horizontal red line). The horizontal black dashed line is for reference showing perfectly accurate estimation. The numerical experiments are repeated with the constraint that promoter switching is slow and the results are shown by box-whisker plots in the right column. The plots show that the source of extrinsic noise and timescale separation determine the size and sign of estimation bias. See main text for details.

To answer this question one must precisely define what is meant by fitting the data generated by the telegraph model with extrinsic noise by the standard telegraph model. For the time being, we shall define a matching of the two models to mean that we have found a set of parameter values of the standard telegraph model for which the first three moments of the mRNA counts of this model matches the first three moments of the mRNA counts of the telegraph model with extrinsic noise. Note that we cannot match fewer or more moments than three since we need to solve three equations to exactly determine the values of three unknowns. Let the *i*th factorial moments of the mRNA counts as given by the standard telegraph model be 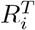 and those of the telegraph model with extrinsic noise be R_*i*_; then the matching procedure is equivalent to setting 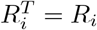 for i = 1, 2, 3. Since 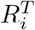 are functions of the telegraph model parameters {α, β, θ}, we have three simultaneous equations that can solved for these parameters, leading to

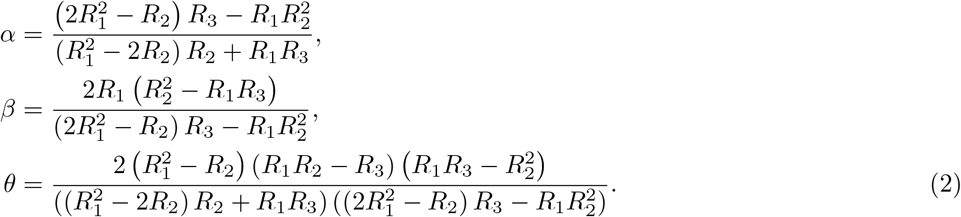

Exact expressions for R_*i*_ for each of the three cases of extrinsic noise (in the synthesis rate, the switching rate or the switching off rate) are derived in Appendix A. Another parameter of interest is the burst size defined as λ = α/θ which is equal to the mean number of transcripts produced while the promoter is in the active state.

To understand systematic bias introduced by the neglection of extrinsic noise in the inference framework, we devise the following procedure: (i) we generated a million random parameter sets with the constraints that the coefficient of variation (CV defined as the standard deviation divided by the mean) of the gamma distribution is less than 1 and the mean of the three transcriptional parameters is in the range 0.1 −200 (values normalised by the mRNA degradation rate). The range of these rates covers the transcriptional parameters estimated by previous studies for mouse and human fibroblasts [7]; (ii) for each parameter set, we inferred the effective parameters using Eq. (2). We discarded parameter estimates which were negative since these are non-physical. The statistics of the positive inferred parameters (normalised by the ground truth values) are shown by box-whisker plots in the middle column of Fig. 2.

The ground truth values are the population averaged parameters in the telegraph model with extrinsic noise. We shall use the angled brackets ⟨⟩ to denote the average over a distribution of parameters. For noise in the synthesis rates, the ground truth values are the mean synthesis rate ⟨*ρ*⟩ and the mean burst size ⟨*b*⟩ = ⟨*ρ*⟩/*k*_off_. For noise in the switching on rate, the ground truth value is the mean switching on rate ⟨*k*_on_⟩. For noise in the switching off rate, the ground truth values are the mean switching off rate ⟨*k*_*off*_⟩ and the mean burst size 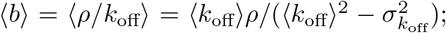 the latter result was derived by integrating the burst size over a gamma distribution with mean ⟨*k*_off_⟩ and variance 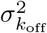 (under the assumption that 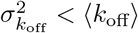 as required by one of our constraints above).

The main conclusions of our numerical experiments are as follows. Extrinsic noise in the synthesis rate causes an overestimation of the synthesis rate and switching rates but an underestimation of the average burst size (Fig. 2a). Extrinsic noise in the switching on rate leads to the opposite (Fig. 2b). Extrinsic noise in the switching off rate leads to an overestimation of the synthesis rate and the burst size and an underestimation of the switching rates (Fig. 2c). These results show that depending on the source of extrinsic noise, there is a specific “bias signature”. Clearly the noise in one parameter causes estimation bias not only in that parameter but in all the others as well. Another difference distinguishing the telegraph model with three different sources of extrinsic noise is the fraction of estimated parameter sets that are positive: merely 0.7% for extrinsic noise in the synthesis rate, 100% for extrinsic noise in the switching on rate and 82% for extrinsic noise in the switching off rate. We note that negative estimated parameter values occur when it is not possible to match the first three moments of mRNA counts of the telegraph model with extrinsic noise with those of the standard telegraph model – in these cases, typically it is possible to only match a smaller number of moments (we study this issue in more depth later).

We repeated the numerical experiment but for the case of slow promoter switching where the synthesis rate is in the range 0.1 −200 while the switching on and off rates are in the range 0.1 −1. The estimated rates as ratios of the ground truth parameters are shown by box-whisker plots in the right column of Fig. 2. By comparison to the previous numerical experiments where all the rates were in the range 0.1 −200, it is clear that slow promoter switching has little effect on estimation bias when extrinsic noise is in the synthesis rate but it significantly diminishes estimation bias when extrinsic noise is in the switching rates; in the latter cases the sign of the bias is unchanged except for the burst size which was overestimated previously and now the reverse occurs. In the slow promoter switching regime, inaccuracy in parameter estimation principally stems from extrinsic noise in the synthesis rates. Interestingly we also find a change in the fraction of parameter sets leading to positive estimates: 86%, 100% and 100% for extrinsic noise in the synthesis, switching on and switching off rates, respectively.

Our inference procedure is based on precise moment-matching between models with extrinsic and without extrinsic noise. Previous studies have noted that for certain models of gene expression, matching of the moments to those of experimental data gives different results than matching of the mRNA count distributions [39]. However we find this is not the case for our inference procedure: matching of the first three moments of the standard telegraph model to those of the telegraph model with extrinsic noise, when it resulted in positive parameter estimates, also led to a good match between the distributions of the two types of models. In Fig. 3(a)-(c) we show three specific examples of the excellent fits, one for each source of extrinsic noise. These are not isolated cases – distribution matching over large swathes of parameters space is confirmed by the high linearity of the P-P probability plots in Fig. 3(d)-(f). Hence we conclude that the results shown in Fig. 2 mirror those that would be obtained by the much more laborious process of matching the distributions of mRNA counts of the standard telegraph model with that from the telegraph model with extrinsic noise. Incidentally this suggests that a model selection algorithm would likely favour the telegraph model since this model has the smaller number of parameters and yet matches the count distributions as well as the extrinsic noise model used to generate the data.

**FIG. 3.**
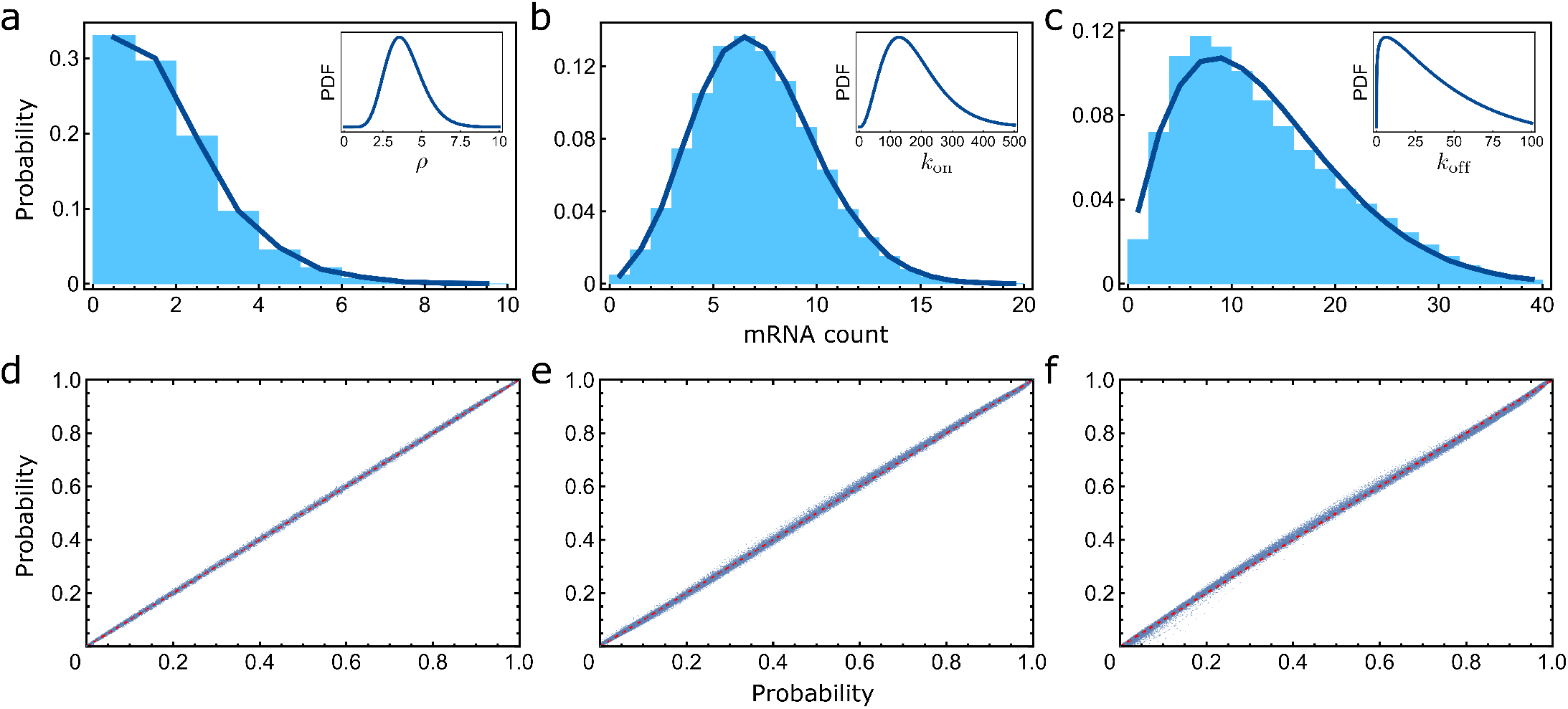
The steady-state mRNA count distributions of the telegraph model with extrinsic noise and the standard telegraph model are well matched, even when the inferred parameters are very different from the true parameters. (a,b,c) compare the distributions with extrinsic noise (histograms) with those of the fitted telegraph model (solid blue line) where the parameters of the latter are estimated using three-moment-matching procedure given by Eq. (2). The distributions of the parameters that are varying from cell-to-cell are shown in the insets. See Appendix C for the values of the inferred parameters and of the true population averaged parameters. Distribution matching is further confirmed by the high linearity of the P-P probability plots (cumulative distribution function for the telegraph model with extrinsic noise vs cumulative distribution function for the standard telegraph model with parameters from three-moment-matching) for 1000 of the parameter sets used for the plots in the middle column of Fig. 2 for variable synthesis rate (d), variable switching on rate (e) and variable switching off rate (f). Similar results are found for parameter sets estimated under slow promoter switching conditions (right column of Fig. 2).

The results derived thus far assume that the distribution of parameters across cells is well approximated by a gamma distribution. In Appendix B we derive results for any distribution of positive-valued parameters provided extrinsic noise is small, i.e. the coefficient of variation of the parameter distributions is small. Surprisingly, we find that when it can be analytically determined, the sign of the estimation bias is the same as in Fig. 2 thus verifying the presence of a universal “bias signature” depending on which rate is the most variable amongst cells.

## III. MASSIVE OVERESTIMATION OF PARAMETERS NEAR A CRITICAL EXTRINSIC NOISE THRESHOLD

We now study in detail the case where the gamma-distributed extrinsic noise is present in the synthesis rate. This case is particularly interesting because: (i) it is the only case where Eq. (2) leads to equations for the inferred parameters that are amenable to analytical analysis (the other two cases are in terms of the exponential integral); (ii) this type of noise is the only one which leads to significant estimation bias independent of the speed of promoter switching (Fig. 2); (iii) if promoter switching is not slow, this type of noise is the only one which leads to many instances where the inferred parameters are negative and hence physically meaningless; (iv) the dependence of the synthesis rate on the cell volume has recently been shown to be a major source of extrinsic noise for many eukaryotic genes [14, 15, 19].

Using the expressions derived for the factorial moments of mRNA counts for gamma extrinsic noise in the synthesis rate with mean ⟨*ρ*⟩ and variance 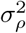 in Appendix A, we find that Eq. (2) simplifies to

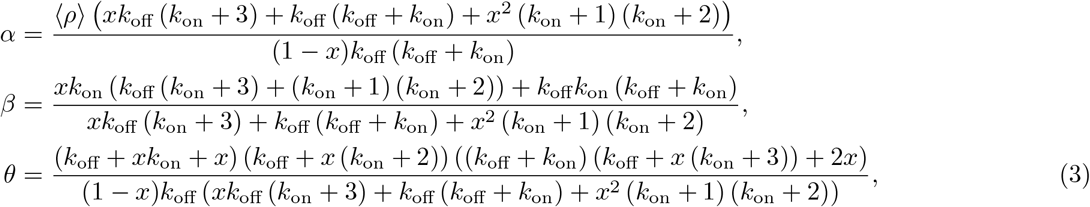

where *x* is the normalised coefficient of variation squared of the extrinsic noise in the synthesis rate defined as

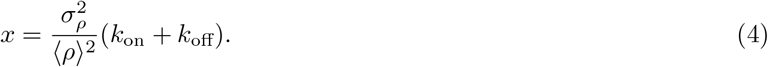

There are several observations that can be made.

1. The inferred mean synthesis rate α is only positive if the size of extrinsic noise is below the critical threshold of *x* = 1. In this parameter regime, the inferred synthesis rate is always larger than the true rate (*α* ≥⟨*ρ*⟩) and it increases monotonically with the size of extrinsic noise (*dα*/*dx* > 0) tending to ∞ as (1 − *x*)^*−*1^.
2. The inferred mean switching on rate *β* is positive, independent of the size of extrinsic noise. If the noise is below the critical value of *x* = 1, the inferred rate is always larger than the true rate (*β* ≥*k*_on_). The estimation error is zero (*β* = *k*_on_) when *x* = 0 and *x* = 1. The maximum error is reached when

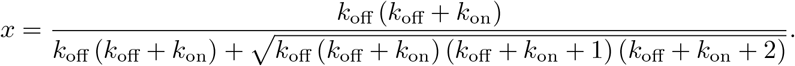
3. The behaviour of the inferred switching off rate *θ* with *x* is similar to that of the inferred mean synthesis rate, i.e. it increases monotonically with *x* tending to *∞* as the critical threshold of *x* = 1 is approached.
4. The inferred burst size (λ = *α/θ*) is positive, independent of the size of extrinsic noise – however note its value is meaningless above the critical threshold because in this case it is the ratio of two negative numbers. If the noise is below the critical value of *x* = 1, the inferred value is smaller than the true value (*λ* < *b* = ⟨*ρ*⟩ /*k*_off_). At the critical extrinsic noise strength of x = 1, the inferred burst size equals ⟨*ρ*⟩ /(*k*_on_ + *k*_off_). When noise is below this critical threshold, the inferred burst size decreases with the size of extrinsic noise if 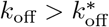 otherwise it reaches a minimum for some value of *x*. The critical value of k_off_ is

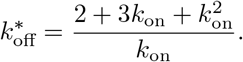

Of particular importance is the finding that near the critical threshold of x = 1, extrinsic noise leads to a very large amplification of the values of the mean synthesis rate and of the mean switching off rate but not of the switching on rate – *this implies that extrinsic noise in the synthesis rate mimics the conditions necessary for transcriptional bursting in the standard telegraph model (without extrinsic noise)* [7].

The results also show that negative parameter values for the synthesis and switching off rates are estimated when noise is larger than the critical threshold of x = 1; given the definition of x in Eq. (4) this means that such non-physical estimates occur when the coefficient of variation squared of extrinsic noise 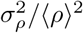 is greater than 1/(*k*_on_ +*k*_off_). Hence it is unlikely that the size of extrinsic noise is large enough to cause such effects when promoter switching is slow whilst it is very plausible if one or both of the switching rates is large – this explains the previous numerical observation in Section II that when the parameter ranges were 0.1 −200, only 0.7% of the parameter sets of the telegraph model with extrinsic noise in the synthesis rate led to positive values of the standard telegraph model, whereas this percentage rose to 86% when the parameter ranges of the switching rates were restricted to 0.1 − 1. We note that while previously it has been reported that estimating parameters by fitting the moments of the standard telegraph model to those of data can lead to negative parameter values, it was not clear why this is the case [40]. One possibility is that this is caused by the finite sample size which leads to errors in moment estimation. However here we have shown that even if the number of cells is infinite (which we simulate by using the exact moments of the telegraph model with extrinsic noise), the inferred parameter estimates from the standard telegraph model can still be negative if the extrinsic noise is larger than the critical threshold.

In Appendix D we show that the estimated parameters show similar qualitative dependence with extrinsic noise strength if the parameter distribution is changed from gamma to lognormal. Since parameter variation between cells is often due to different protein numbers (for e.g. RNA polymerase in the case of synthesis rates) and these are commonly measured to be either gamma or lognormal [41–45] it follows that the results derived here have wide applicability. Interestingly, we find that the critical extrinsic noise strength is smaller for lognormal noise compared to gamma noise, thus suggesting that massive overestimation of the switching off and synthesis rates might be widespread.

### A. Maximum likelihood estimates are in good agreement with the moment-based inference theory

Inference using the standard telegraph model is often done using the method of maximum likelihood [6, 13, 46]. This is a different procedure than three-moment matching and hence the question is whether the estimation bias for x < 1 predicted by our moment-based theory is also reflected in the maximum likelihood estimates. We directly answer this question as follows. For a given extrinsic noise strength (defined by x), we performed 40 simulation experiments, in each of which sampling of the steady-state distribution of the telegraph model with extrinsic noise in the synthesis rate was used to obtain mRNA counts in 10^4^ cells. For each experiment, we infer the rates by maximising the likelihood of the standard telegraph model evaluated using the simulated count data; for details see Appendix E. The statistics of the 40 sets of estimated rates are shown using box-whisker plots in Fig 4(a)-(d). Note that the moment-based theory (blue dashed line) given by Eqs. (3) is close to the median of the rates estimated using maximum likelihood (horizontal red lines). The parameters used for this example are such that the gene spends most of its time in the off state. In Fig. 7(a)-(d) we show the same holds for another parameter set where the gene spends most of its time in the on state – in this case the estimation bias becomes huge – for example the switching off rate and the burst size, as estimated by the standard telegraph model, are about 40 times larger and 5 times smaller respectively than the true values when extrinsic noise is only moderately large x = 0.45. In Fig. 7(e)-(h) we also show that for all strengths of extrinsic noise, the mRNA count distribution of the telegraph model with the inferred parameters cannot be distinguished from the mRNA count distribution of the telegraph model with extrinsic noise – this is in line with the findings using moment-based inference in Fig. 3. In summary our results confirm that the bias predictions of our moment-based theory apply also to the commonly used maximum likelihood method when the number of cells for which mRNA counts are measured is not too small (10^4^).

**FIG. 4.**
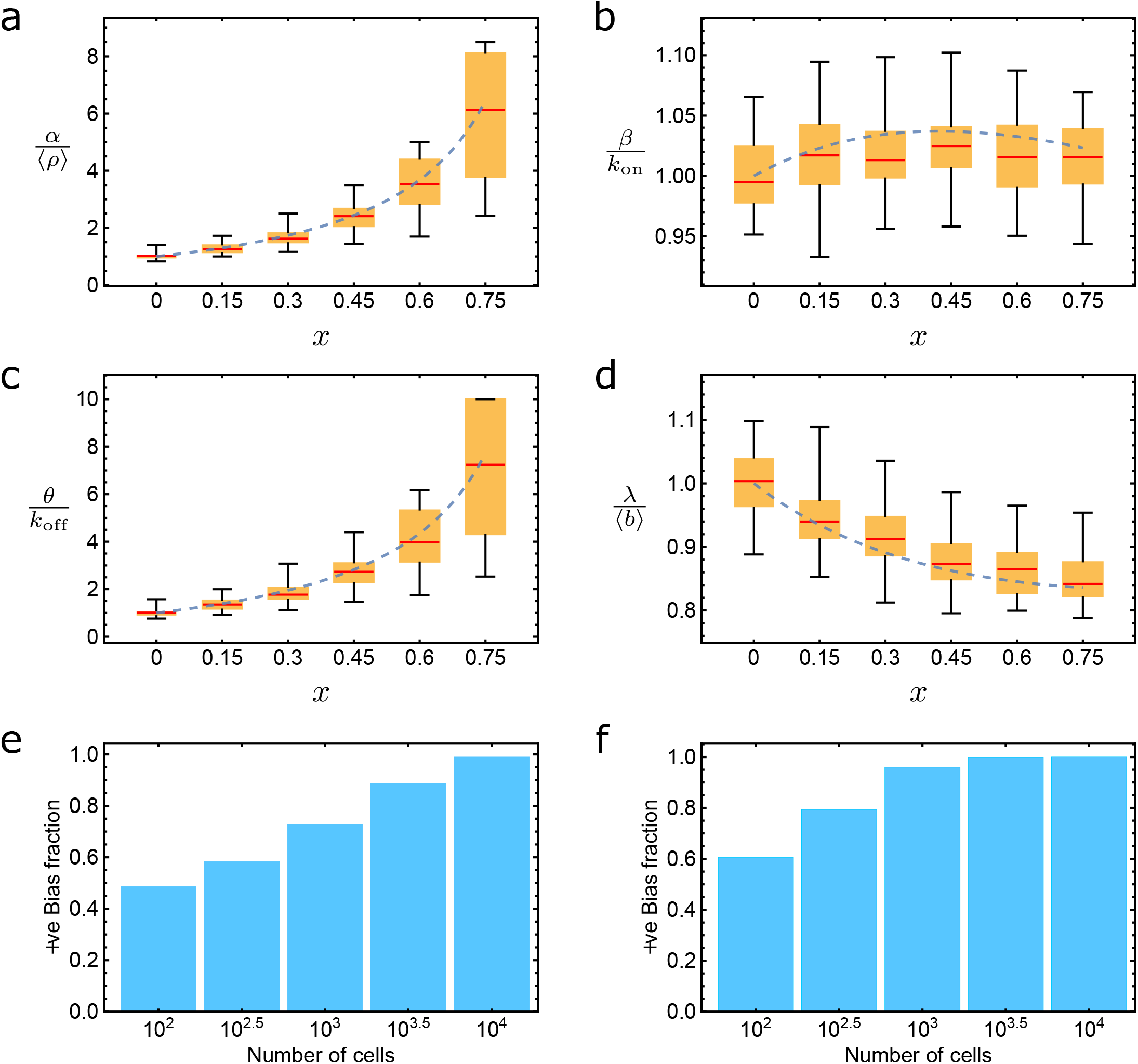
Comparison of the transcriptional parameters estimated using moment-based theory and those estimated by maximum likelihood fitting of the telegraph model to simulated mRNA count data from a population of cells with cell-to-cell variation in the synthesis rate *ρ* (gamma distributed). The rates *α, β, θ, λ* = *α/θ* are the synthesis rate, switching on rate, switching off rate and burst size of the standard telegraph model, respectively. While ⟨*ρ*⟩, *k*_on_, *k*_off_, ⟨*b*⟩ = ⟨*ρ*⟩ */k*_off_ have the same meaning but for the telegraph model with extrinsic noise. The latter are the true parameters with which we compare the parameters estimated using the standard telegraph model. Blue dashed lines show the predictions of the moment-matching theory given by Eq. (3). The box-whisker plots (a-d) summarise the statistics of the maximum likelihood rates inferred from a population of 10^4^ cells; the distribution for each rate is obtained by repeating the inference for 40 samples, each having 10^4^ cells. The lower and upper fences show the min and max, respectively; the yellow bar shows the interquartile range; median is shown by the horizontal red line. The parameters of the telegraph model with extrinsic noise are ⟨*ρ*⟩ = 20, *k*_on_ = 1, *k*_off_ = 5, ⟨*b*⟩ = 4 (gene spends 5*/*6 of its time in the inactive state). The size of extrinsic noise *x* is equal to the normalised coefficient of variation squared as given by Eq. (4). The moment-based theory is close to the median of maximum likelihood estimates. The positive bias fraction (the fraction of parameter estimates for which *α* and *θ* are larger than the ground truth values) is generally a function of the number of cells; this is shown in (e) and (f) for *x* = 0.2 and *x* = 0.6, respectively. Bias (positive bias fraction *>* 0.5) is evident for samples with a number of cells larger than about 10^3^ and even for less when the extrinsic noise is strong. Parameter values are the same as for (a)-(d). See main text for details.

While some smFISH and RNA sequencing experiments collect this much data (or more) [13, 47] others do not [6, 48]. If the sample size is very small then the moments may not be well estimated and hence our theory may not accurately capture the magnitude of estimation bias. Hence we used simulations to investigate the extent of bias in the maximum likelihood estimates of the synthesis and switching off rate (the rates displaying the largest bias in Fig. 4 (a,c)) as a function of sample size. In Fig. 4(e)-(f) we show the results for small (*x* = 0.2) and moderately large (*x* = 0.6) extrinsic noise, respectively, for the same parameters as Fig. 4(a)-(d). For a given number *N* of cells, the positive bias fraction was calculated as follows: we generated 500 samples each with *N* cells, performed parameter inference for each sample using the maximum likelihood method and then calculated the fraction which overestimated *α* and *θ*. When samples had a small number of cells (order of a hundred) and the noise was small, the bias fraction was close to 1/2 indicating that in this case it is difficult to discern any systematic bias since it is overshadowed by the large random variation in parameter estimates from one sample to another. In contrast when the sample size was of the order of a thousand cells or larger, the positive bias becomes evident. Same occurs for large noise but now the (larger) bias is observable even for samples with only order hundred cells.

## IV. ESTIMATION BIAS IN THE PRESENCE OF TRANSCRIPTIONAL BURSTING: OVERESTIMATION OF BURST SIZE AND UNDERESTIMATION OF BURST FREQUENCY

### A. Gamma-distributed burst size

When a gene spends most of its time in the off state, i.e. taking the limit of large k_off_ at constant k_on_, according to Eq. (4) the critical threshold of x = 1 corresponds to a coefficient of variation squared of extrinsic noise that is equal to 0. Hence in this case it is generally impossible to match the first three moments of the mRNA counts as predicted by the standard telegraph model to the moments of the telegraph model with extrinsic noise in the synthesis rate (the estimated switching off and synthesis rates will be negative for any extrinsic noise size). In addition, since in this case x can be very large, it follows from Fig. 4(e) and (f) that positive bias will be evident even in small sample sizes of the order of hundred cells. We note that transcriptional bursting is a particular case of the large k_off_ limit; it only manifests when the burst size is also large enough. As we now show, we can still obtain insight into the effects of extrinsic noise in parameter estimation if we only match the first two moments of mRNA counts to those of a reduced version of the telegraph model.

The simplest two parameter distribution which is most often fitted to mRNA count data, especially that from sequencing, is the negative binomial distribution [49–53]. This mRNA distribution is the steady-state solution of the chemical master equation for the reaction scheme [54]:

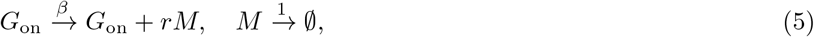

where r is an integer random number sampled from the geometric distribution supported on the set 0, 1, 2, … Specifically this models the case where a gene produces bursts of mRNA with mean burst size λ = ⟨*r*⟩ at a frequency *β*. The steady-state negative binomial distribution solution of this model is

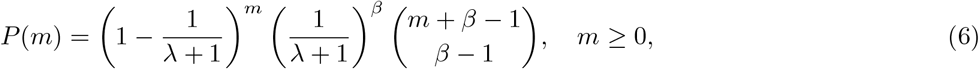

where *m* is the mRNA count per cell. This distribution is completely defined by its first two factorial moments:

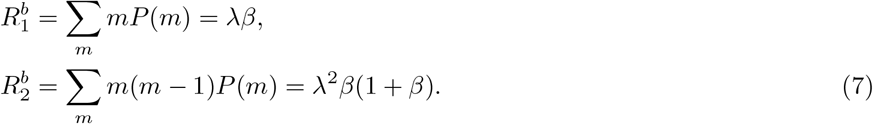

Equating these to the first and second factorial moments of the telegraph model with gamma distributed noise on the synthesis rate (see expressions for *R*_1_ and *R*_2_ in Appendix A), we find the estimated rates are given by:

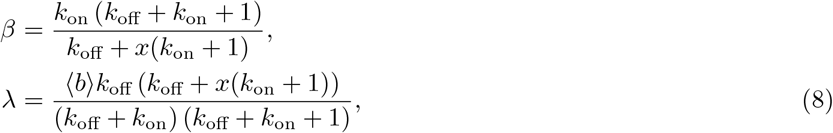

where ⟨*b*⟩ = ⟨*ρ*⟩ /*k*_off_ is the true burst size and *x* is the (normalised) size of extrinsic noise as defined in Eq. (4). Note that a gamma distributed synthesis rate is equivalent to a gamma distributed burst size. The fitting procedure is illustrated in Fig. 5(a). Clearly an advantage of this type of model fitting is that the parameters inferred are always positive, independent of the strength of extrinsic noise. On the other hand, note that the errors made by this type of inference can be significant even when there is no extrinsic noise (*x* = 0). In this case, the burst frequency should be *β* = *k*_on_ (the rate at which a burst starts is equal to the rate at which the gene switches on) while the burst size should be *λ* = ⟨*b*⟩. This is only obtained if the condition *k*_off_ ≫1 + *k*_on_ is true which makes sense when one considers that the steady-state mRNA distribution solution of the standard telegraph model is well approximated by a negative binomial in this limit [7]. Enforcing the condition *k*_off_ ≫ 1 + *k*_on_, we find Eq. (8) reduces to:

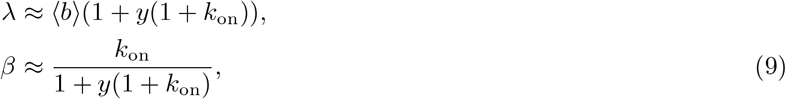

where *y* = *x*/*k*_off_ is to a very good degree of approximation equal to 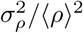, the coefficient of variation of extrinsic noise squared. From these results, it is clear that *fitting a negative binomial distribution to single-cell data characterised by transcriptional bursting and gamma distributed extrinsic noise in the synthesis rate will overestimate the burst size and underestimate the burst frequency*.

**FIG. 5.**
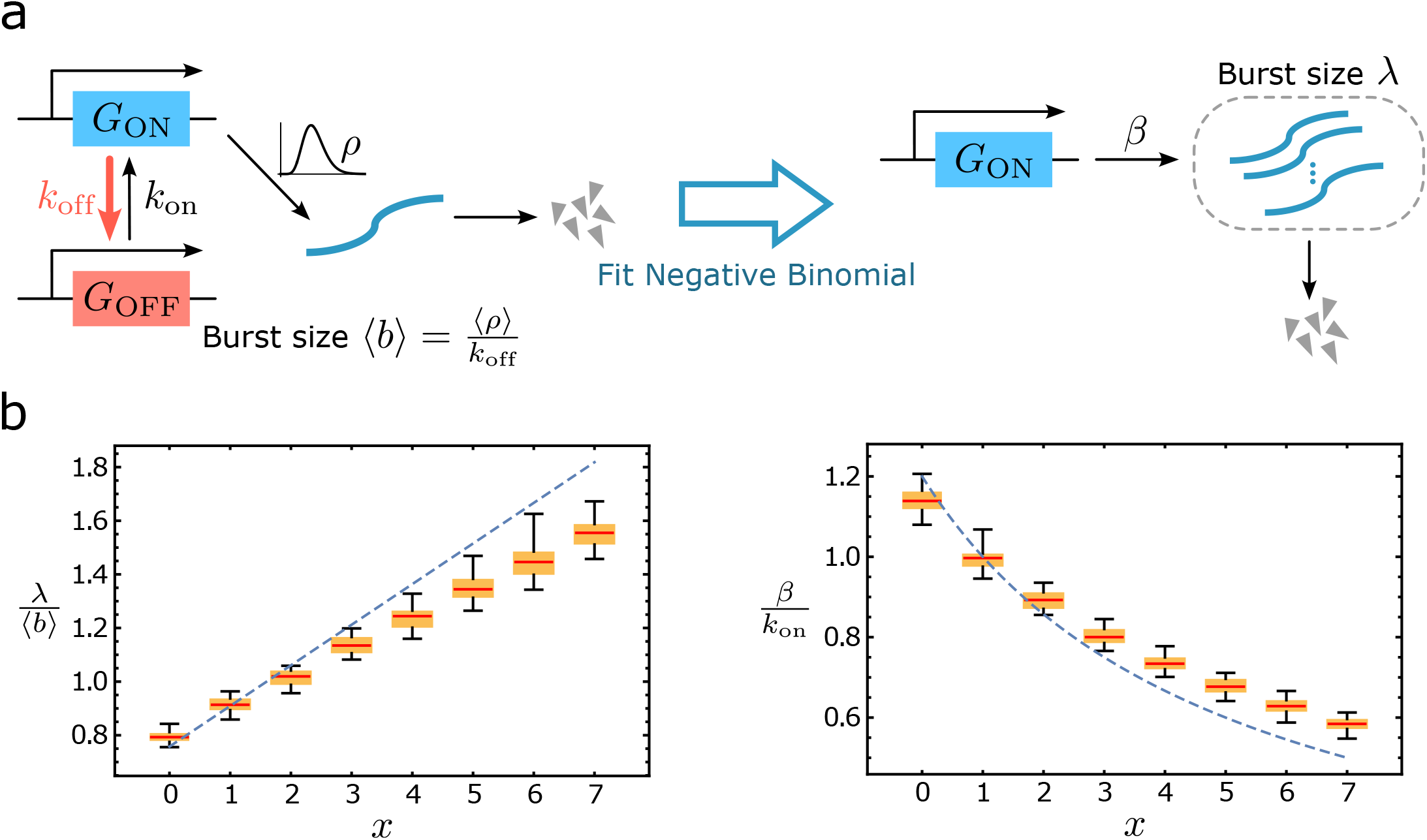
Estimation bias for extrinsic noise in the synthesis rate in the presence of transcriptional bursting. In (*a*) we consider the case where parameters are estimated by fitting a negative binomial distribution using maximum likelihood to simulated mRNA count data in a population of cells with cell-to-cell variation in the synthesis rate (gamma distributed) and transcriptional bursting (large switching off and synthesis rates compared to the switching on rate). The negative binomial distribution used for maximum likelihood fitting is given by Eq. (6). In (*b*) the box-whisker plots (min, max, interquartile range and median) summarise the statistics of the inferred rates using maximum likelihood from 40 simulation experiments, in each of which there are 10^4^ cells. The moment-based theory (dashed blue lines) is given by Eq. (8). The trend of maximum likelihood estimates are well described by the moment-based theory – the presence of extrinsic noise causes the burst size to be overestimated and the burst frequency to be underestimated. However as the size of extrinsic noise *x* increases, there are systematic deviations which derive from deviations of the distribution of mRNA counts from a negative binomial. The parameters of the telegraph model with extrinsic noise are ⟨*ρ*⟩ = 20, *k*_on_ = 1, *k*_off_ = 10, ⟨*b*⟩ = 2 (gene spends 10*/*11 of its time in the inactive state). The size of extrinsic noise *x* is the normalised coefficient of variation squared defined by Eq. (4).

In Fig. 5(b) we compare the predictions of the moment-based theory (dashed blue lines) given by Eqs. (8) with estimates obtained by fitting a negative binomial distribution using maximum likelihood (shown using box-whisker plots) to simulated data for 10^4^ cells. There are two main observations: (i) for *x* = 0, the burst size and burst frequencies are not equal to the true parameters. The reason is because the formal condition *k*_off_ ≫1 + k_on_ necessary for the telegraph model’s solution to be well approximated by a negative binomial distribution is only approximately met for the parameters used in this example (*k*_off_ = 10, *k*_on_ = 1). This shows that even in the absence of extrinsic noise, there are typically systematic errors in parameter estimation (underestimation of burst size and overestimation of burst frequency). These differences approach zero as the switching off rate is made to be 100 times or larger than the switching on rate; (ii) the maximum likelihood estimates follow the trends predicted by the moment-based theory, namely that an increase in extrinsic noise leads to an amplification in the observed burst size and an attenuation of the observed burst frequency. However the differences between the two methods increase with the size of extrinsic noise. This is not a sample size effect – repeating the inference using 100 times more cells (40 experiments each with 10^6^ cells) does not remove the differences. We note that the negative binomial is a two parameter distribution that can be estimated completely from knowledge of its two moments, i.e. if the distribution of the telegraph model with extrinsic noise is well approximated by the negative binomial then the moment-based theory and maximum likelihood would be in good agreement. Hence the differences between the two methods, with increasing extrinsic noise size, ensue from the lack of “flexibility” of the negative binomial distribution to describe the distribution of mRNA counts.

### B. General case: extrinsic noise in burst size and frequency with arbitrary distributions

We next consider the general case where there is transcriptional bursting and arbitrarily distributed noise on any of the three parameters of the telegraph model (synthesis rate or one of the switching rates). Since we have bursting rather than using the full telegraph model with extrinsic noise, instead we consider the following reaction system as a suitable reduced model describing the data:

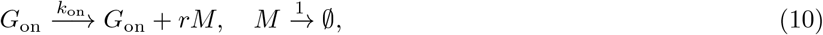

where ⟨r⟩ is the mean burst size b and k_on_ is the burst frequency. The cell-to-cell variations in these two parameters are described by a burst frequency distribution, f(k_on_) and a burst size distribution, g(b). It follows that the first two factorial moments of the mRNA counts are given by integrating the factorial moments of the negative binomial distribution solution of the model in the absence of extrinsic noise over the distribution of parameters:

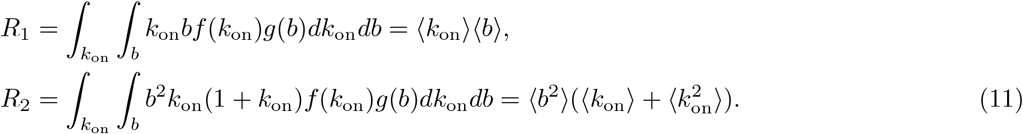

We simulate the fitting of the data to a non-extrinsic noise model (Fig. 5a) by setting 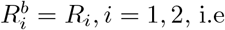. equating Eq. (7) with Eq. (11). Then solving for the estimated mean burst size and the mean burst frequency we find:

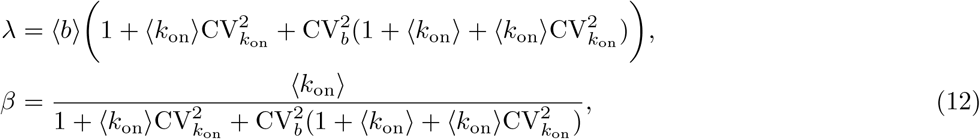

where 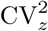 is the coefficient of variation squared of parameter z where z = k_on_, b. In the limit of no extrinsic noise in the switching on rate (no extrinsic noise in the burst frequency), Eq. (12) reduces to Eq. (9). Note that Eq. (12) has a powerful interpretation: *if transcriptional bursting is present, independent of the source of extrinsic noise and the inherent distribution of the parameters, the mean burst size is overestimated while the mean burst frequency is underestimated when these are deduced from fitting a negative binomial distribution (or the mRNA count distribution of the standard telegraph model) to the mRNA count data*. The theory’s accuracy is tested and confirmed using maximum likelihood applied to simulated counts data generated using the telegraph model with gamma distributed extrinsic noise for four parameter sets (two with noise in the burst frequency and two with noise in the burst size) in Table I.

**TABLE I.** Comparison of maximum likelihood estimation using a negative binomial distribution and Eq. (12). For each set of parameters, we generated 100 independent samples, each with 10^4^ cells. The mRNA counts per cell were simulated from the telegraph model with gamma distributed extrinsic noise in the switching on rate (lines 1 and 3) or in the synthesis rates (lines 2 and 4). For each sample we estimated the burst frequency and size by fitting a negative binomial distribution using maximum likelihood. The last two columns show the estimated parameters ([first quartile,median,third quartile]) computed over all 100 samples. The moment-based theory predicts that in all these cases, *λ/* ⟨*b*⟩ *≈*11 and *β/* ⟨*k*_on_⟩ *≈*1*/*11. The median of the maximum likelihood estimates varies little between the four parameter sets and its value is close to the theoretical predictions.

| $\langle \rho \rangle$ | $\langle k_{\text{on}} \rangle$ | $k_{\text{off}}$ | Noise | $\lambda/\langle b \rangle$ | $\beta/\langle k_{\text{on}} \rangle$ |
| --- | --- | --- | --- | --- | --- |
| 600 | 10 | 300 | $CV_{k_{\text{on}}} = 1$ | [10.45, 10.55, 10.68] | [0.088, 0.089, 0.090] |
| 200 | 10 | 200 | $CV_b = 1$ | [10.30, 10.46, 10.62] | [0.090, 0.091, 0.092] |
| 400 | 10 | 600 | $CV_{k_{\text{on}}} = 1$ | [10.59, 10.75, 10.87] | [0.090, 0.091, 0.092] |
| 800 | 10 | 500 | $CV_b = 1$ | [10.53, 10.70, 10.82] | [0.091, 0.092, 0.093] |

### C. Application to scRNA-seq data

A recent study has used the method of maximum likelihood to fit the mRNA counts from allele-sensitive single-cell RNA sequencing to the standard telegraph model thus determining the transcription parameters for endogenous mouse genes transcriptome-wide [6]. For a large number of genes, the switching on rate was found to be much less than the switching off rate and the synthesis rate thus leading to the conclusion that transcriptional bursting is ubiquitous in mammalian genes. This suggests that the distributions of mRNA counts for many genes in this dataset are well fit by a negative binomial distribution, i.e. the only two parameters that can be determined accurately are the burst frequency (switching on rate) and the burst size (the ratio of the synthesis and switching off rates). In fact, as we show in Fig. 8, to a good degree of approximation their standard telegraph model estimates (using the method of maximum likelihood) for the burst frequency and size for genes displaying highly bursty expression are in good agreement with those obtained by simply equating the first and second factorial moments of the mRNA count data for each gene to the factorial moments Eq. (7) predicted by the bursty expression model (5) and solving for λ and β. Any deviations between the two estimates are due to the small sample size of a few hundreds of cells.

**FIG. 6.**
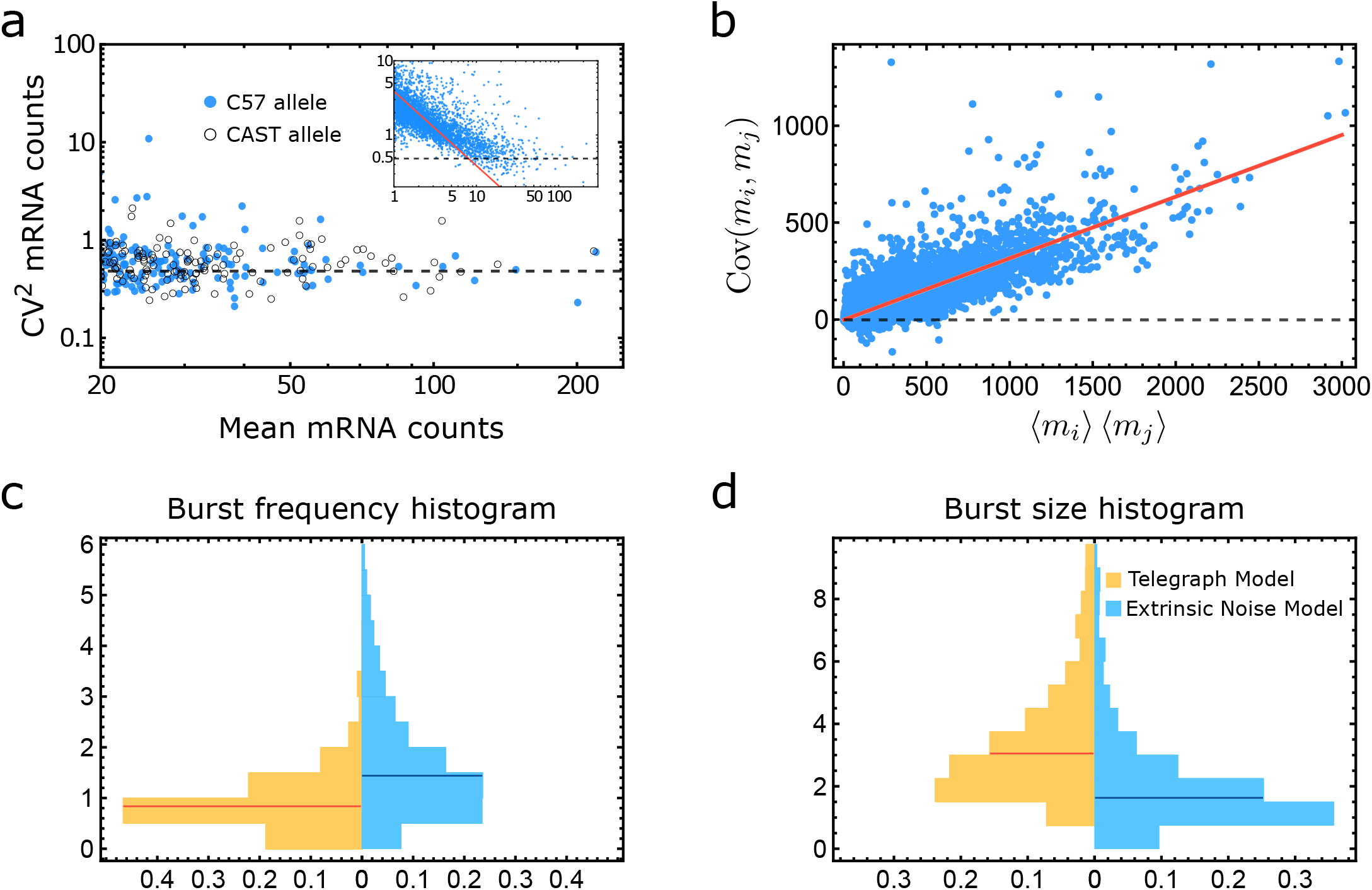
Estimation bias of burst size and frequency from scRNA-seq data [6]. (a) Plots of the coefficient of variation squared CV^2^ of mRNA counts versus their mean for the C57 and cast alleles of high expression genes (defined as those with mean *>* 20) in 224 primary adult fibroblasts – each dot is a gene. The CV^2^ does not vary much with the mean expression (dashed black line) indicating a high degree of extrinsic noise from a cell-specific factor. In contrast for low-expression genes, there is a fuzzy but clear inverse relationship between CV^2^ and the mean number of counts (solid red line in the inset), indicating strong contributions from both intrinsic and extrinsic noise. (b) Plot of the covariance between a gene pair (gene *i* and gene *j*, where *i* ≠ *j*) versus the product of their mean expression for 1877 highly bursty genes (C57 allele) (see Fig. 8 for their definition). The plot confirms the theoretical prediction Eq. (15); the red regression line has correlation coefficient = 0.90 and its slope (0.32) is equal to the coefficient of variation squared of extrinsic noise. 96% of gene pairs have a positive covariance. (c) and (d) Paired histograms for the estimates of the burst frequency and the burst size, obtained by matching the first two moments of the C57 data for each of the highly bursty genes to those of the standard telegraph model assuming transcriptional bursting and separately by correcting it to account for extrinsic noise with size as found in (b). The horizontal lines show the medians of the distributions.

**FIG. 7.**
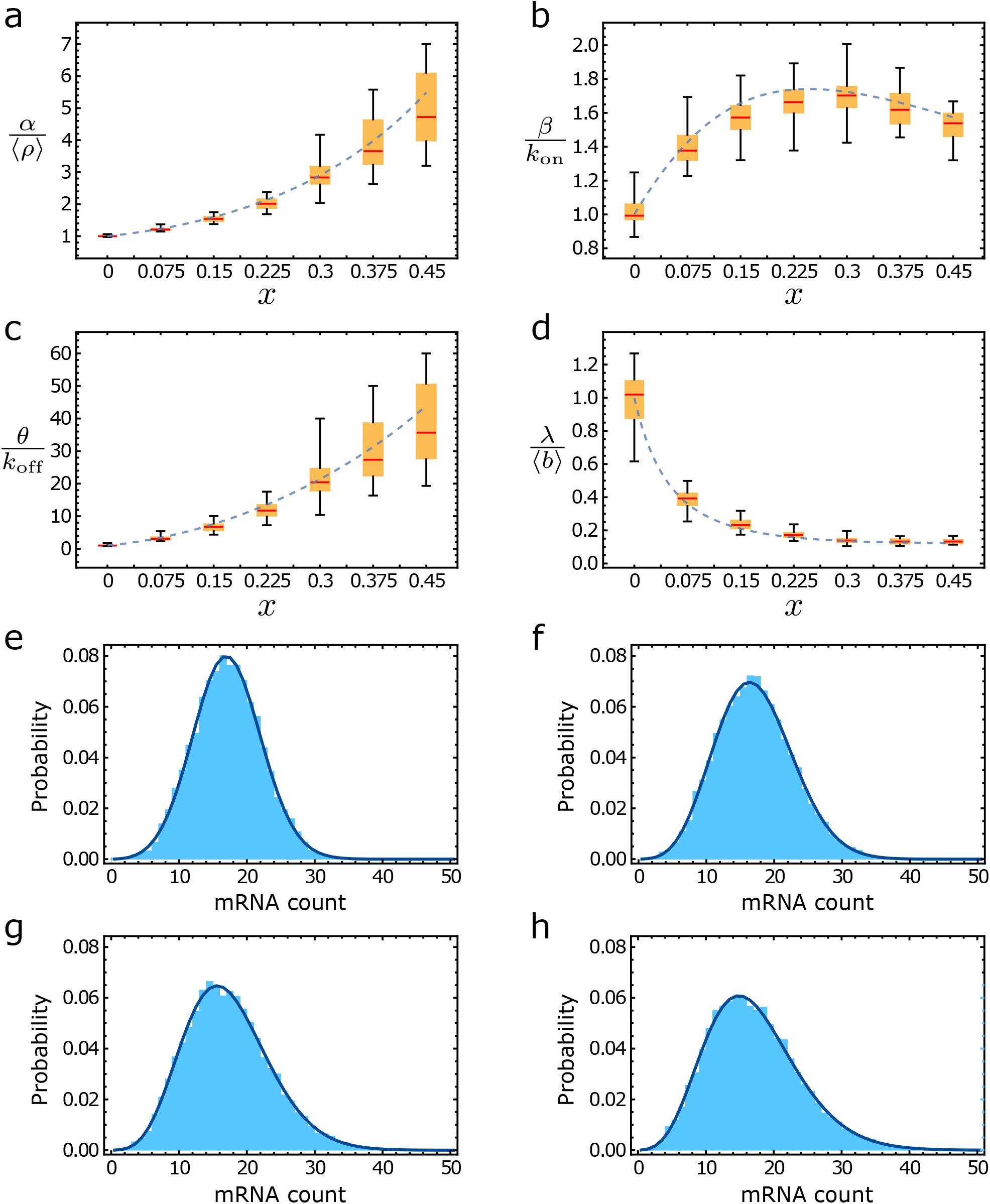
Comparison of the transcriptional parameters estimated using moment-based theory and maximum likelihood fitting of the telegraph model to simulated mRNA count data with a cell-to-cell variation in the synthesis rate *ρ* (gamma distributed) and where the gene spends most of its time in the on state. (a)-(d) shows box-whisker plots similar to those in Fig. 4(a)-(d) with the difference that the parameters of the telegraph model with extrinsic noise are ⟨*ρ*⟩ = 20, *k*_on_ = 5, *k*_off_ = 1, ⟨*b*⟩ = 4 (gene spends 5*/*6 of its time in the active state). In (e)-(h) we show that the corresponding distributions of the telegraph model fitted by maximum likelihood (solid lines) to the simulated data (histogram) for the cases *x* = 0, 0.15, 0.3, 0.45 respectively – clearly, the telegraph model cannot be distinguished from the telegraph model with extrinsic noise from fitting of the population snapshot distributions of mRNA counts per cell.

**FIG. 8.**
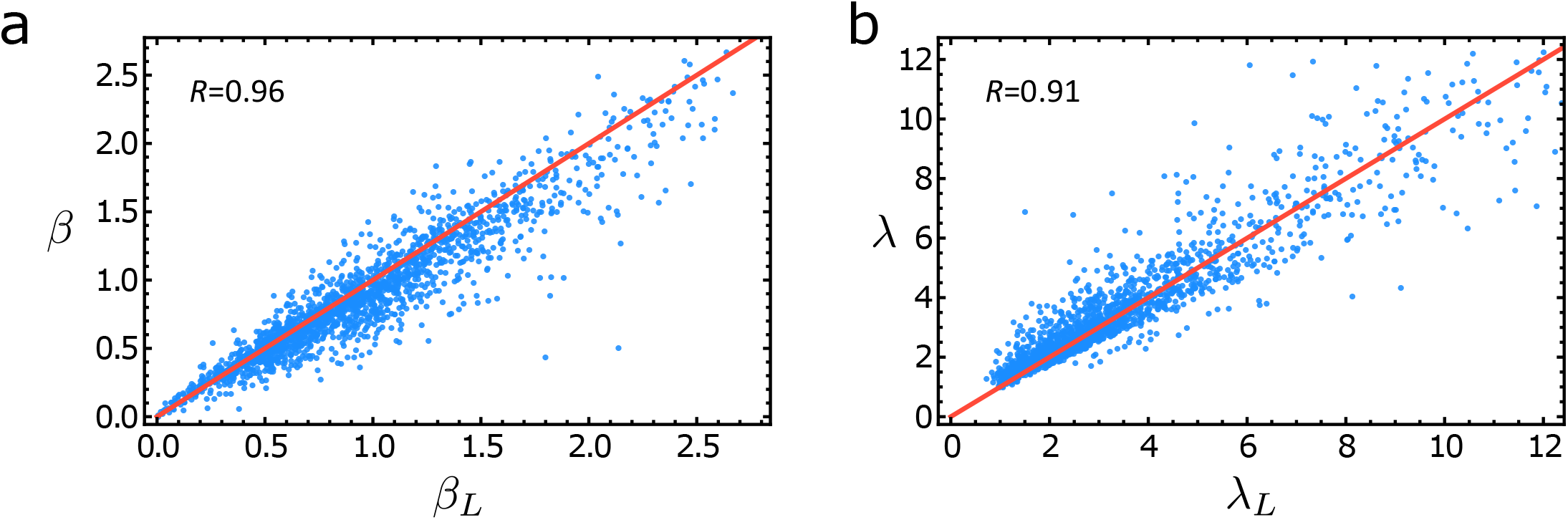
Plots of the burst frequency (a) and size (b) for 1877 bursty genes estimated by fitting sc-RNA seq data from 224 primary adult mouse fibroblasts (C57 allele) to the standard telegraph model using the method of maximum likelihood (x-axis) and separately estimated using two moment-matching (y-axis). The maximum likelihood estimates are found in the file 41586 2018 836 *MOESM* 3 *ESM*.*xlsx* associated with Supplementary Table I in [6] and they are denoted by the subscript *L*. The moment-matching estimates are obtained by equating the first and second moments of the mRNA count data for each gene to the moments Eq. (7) predicted by the bursty expression model (5) and solving for *λ* and *β*. We consider only genes for which the mean expression *>* 1 and the maximum likelihood estimates of switching off rate is at least 20 times larger than the estimate of the switching off rate, thus enforcing transcriptional bursting. The maximum likelihood and moment-matching estimates are similar as indicated by the Pearson correlation coefficient *R* and the closeness of the points to the red lines *β* = *β*_*L*_ and *λ* = *λ*_*L*_.

However since in the transcriptional bursting regime the normalised coefficient of variation of extrinsic noise (Eq. (4)) is large, given the results in Fig. 4(f), we expect that the moment estimates computed from the few hundred cells measured in [6] will be significantly biased due to extrinsic noise. To correct the moment-matching estimates using the theory we previously developed, we need to estimate the coefficient of variation of extrinsic noise. We assume that the data can be well described by the effective reaction scheme (10) with arbitrary distributions of the burst size and burst frequency. Using Eq. (11) it can be shown that the coefficient of variation squared 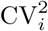 of the mRNA counts for gene i can be written as:

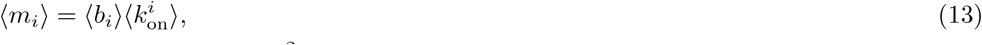

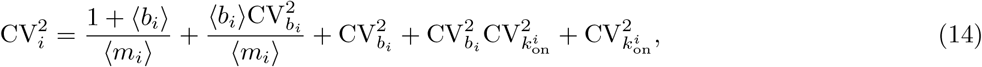

where ⟨m_*i*_⟩ is the mean number of mRNA counts for gene i, and 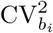 and 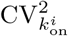 are the coefficient of variation squared for the burst size and frequency of gene i, respectively. The means and coefficients of variation of the parameters are calculated for a population of cells. We find that this law is reflected in the sc-RNA seq data (it has also been found in other scRNA-seq datasets [48, 53]). In Fig. 6(a) we plot the coefficient of variation squared as a function of the mean expression for about 10^4^ mouse fibroblast genes with mean expression > 1. For both the C57 and the CAST alleles, the law 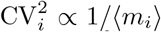 manifests for mean expression below 10 (red line in inset of Fig. 6(a)) followed by a transition to the law 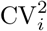 equals some constant (independent of the gene index i) for larger mean expression levels. These observations follow Eq. (14) – the first two terms describe the small expression law and the last three terms explain the high expression law.

The significant scatter about the red line for low mean expression is expected because each gene will have a different value of the mean burst size and hence the first two terms in Eq. (14) can only be approximately inversely proportional to the mean expression. On the other hand, the low scatter about the line CV^2^ equals some constant (for high-expression genes) indicates that the coefficient of variation of the burst size and frequency are similar across genes; these measures of the size of extrinsic noise are also independent of the allele. These observations about extrinsic noise can be explained as follows. Since burst frequency changes with cell cycle phase [4, 13, 14], it is expected that its coefficient of variation is not particularly large for this dataset because the population of cells is mostly in the G1 phase [6]. Thus we assume that the variation in the parameters is dominated by the burst size since this is often influenced by the cell volume [14]. Specifically we assume that the burst size b_*i*_ of gene i in a cell of volume V is given by *b*_*i*_ = *r*_*i*_*V* where r_*i*_ is some gene specific constant. This law naturally emerges if there is concentration homeostasis, i.e. transcript numbers scale approximately linearly with the cell volume thus maintaining a constant concentration [55–57]. Then it immediately follows that 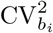 in Eq. (14) equals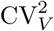, i.e. the coefficient of variation in the burst size is the same for each gene and equal to the coefficient of variation of the cell volume (computed for a population of cells). Note that a rough estimate of the size of extrinsic noise is given by the coefficient of variation squared of mRNA counts in the limit of large mean expression as shown by the dashed black line in Fig. 6(a).

An analysis of the covariance between different genes provides strong supporting evidence of extrinsic noise due to a cell-specific factor (such as the volume). In Appendix F we analytically show that if the synthesis rate (and hence the burst size) linearly scale with a cell-specific factor V then the covariance between the mRNA counts of any two genes i and j (i ≠ j) is given by

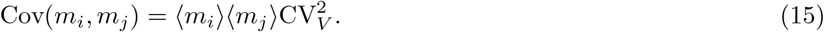

This relationship is true also if the burst frequency has a linear dependence on a cell-specific factor provided some additional criteria are met. Hence the prediction is that extrinsic noise induces a positive covariance between the counts of any pair of genes (see also [58] where high gene-gene correlations where observed and explained by burst size coordination). This might be surprising since one would assume that correlations between some genes would be positive/negative due to regulatory effects and many would be uncorrelated. However calculation of the covariance for highly bursty genes shows that most show a positive covariance and the linear relationship predicted by theory Eq. (15) is evident (Fig. 6(b)). Additionally the slope of the best straight line through the data gives us a reliable estimate of the size of extrinsic noise: 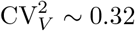

Based on the foregoing arguments we shall assume that for each mouse fibroblast gene displaying apparent bursting behaviour in the sc-RNA seq dataset, the mean expression and the coefficient of variation squared of the mRNA counts are given by Eqs. (13)-(14) with 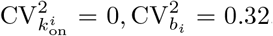. Note that the genes selected for inference are the subset of the highly bursty genes in Fig. 8 for which 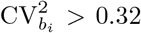. Solving the equations for the mean expression and the coefficient of variation squared simultaneously we obtain estimates of the mean burst size and frequency for each gene. The distribution of these two parameters across the transcriptome are shown in blue in Fig. 6(c) and (d); for comparison, we also show (in yellow) the corresponding distributions if we had to ignore extrinsic noise, i.e. as done in the original analysis of [6]. Clearly, the addition of extrinsic noise leads to an increase in the burst frequency estimates (the mass of the distribution spreads towards higher values) and a corresponding decrease in the burst size estimate (the mass of the distribution concentrates in the small value region) – this is in line with our general theory summarised by Eq. (12). In particular, the median burst frequency increased by about 2 times while the median burst size decreased by the same amount when extrinsic noise was taken into account. We note that what we have referred to as the mean burst size for gene i is the observed mean burst size, i.e. ⟨b_*i*_⟩ = r_*i*_ ⟨V⟩, where the average is taken over the population of cells. If one is interested in deducing r_*i*_ then a measurement or estimation of the average cell volume is additionally needed. The difficulty here is that it is likely that what we have called cell volume is not simply that but a variable which depends on various cell-specific factors and also technical variability [59].

In our inference procedure we have made a major assumption: the number of mRNA per gene per cell corresponds to the number of UMIs (unique molecular identifiers) detected. This is the same assumption made in [6]. Comparing their measured distributions of mRNA counts for 5 genes measured using both smFISH and scRNA-seq (Extended Fig. 7), it is clear that the latter display an accumulation of probability mass to the left implying there is a non-negligible fraction of transcripts in each cell that is not captured by scRNA-seq, a well-known property [60]. It is however not clear how to correct for these effects. Hence the burst frequency and size histograms output by our extrinsic noise moment-matching procedure shown in Fig. 6(c) and (d) are not to be interpreted as the true histograms but rather a step in that direction.

Heuristic methods to correct estimates for capture efficiency of scRNA-seq technology have been recently proposed in [61]. The main idea of the moment-based method proposed (referred to as the MME) is that instead of performing parameter inference directly on raw mRNA counts from UMI count-based scRNA-seq data, one should first normalise the counts from each cell by a heuristic cell-specific scaling factor (calculated from the raw data), and subsequently the moments of these renormalised counts can be fitted to those of a standard telegraph model to estimate the transcriptional parameters. Despite its simplicity and ease of computation, it can be analytically shown that this procedure is flawed because the normalisation of count data does not transform the moments of the telegraph model with extrinsic noise to those of the standard telegraph model. We show in Appendix G that inference using this method is only accurate when the mean expression is large enough – in contrast the inference method whose results are shown in Fig. 6(c) and (d) does not suffer from such inaccuracies. A neural-network (NN) method is also developed in [61] to correct for capture efficiency and this maybe useful when it can be ascertained that the major sources of extrinsic noise are in the synthesis rate, e.g. when the scRNA-data is collected from cells in a particular cell cycle phase such that burst frequency variations between cells are minimised [4, 13, 14]. This method also predicts corrected estimates for the burst size and frequency of the same scRNA-seq dataset we analysed in Fig. 6. The burst size is defined differently than in our case (in our notation for gene i it is defined as the parameter r_*i*_) and hence it is not possible to compare our estimate with those from the NN method. The burst frequencies can however be compared: both the NN method and ours predict that taking into account extrinsic noise considerably widens the burst frequency histogram. Our theory additionally predicts that independent of the source of the extrinsic noise, if transcriptional bursting is evident, the median burst frequency of bursty genes should be larger after correcting for it; this might be consistent with the results of the NN study but it is difficult to ascertain from the published data analysis.

## V. SUMMARY AND DISCUSSION

In this paper, we developed a theory that explains the size and sign of estimation bias when the standard telegraph model is used to infer transcriptional parameters from single cell data. We note that while it is widely appreciated that this model does not take into account extrinsic noise (it assumes the parameters do not vary between cells which is of course unrealistic), it is still the approach of choice in most studies [3, 5, 6, 40, 62] because of its simplicity and the considerable difficulty in completely correcting for all sources of extrinsic noise in mRNA count data. Hence a main motivation for our study was to understand how well the parameters estimated using the standard telegraph model approximate the true average parameter values in a population, i.e. does the concept of a virtual “mean cell” make sense? [26]

A key assumption of our theory is that single-cell data is accurately approximated by the telegraph model with extrinsic noise in either the synthesis rate or the switching on rate or the switching off rate. By matching the moments of this model with those of the standard telegraph model, we mimic the common procedure of fitting the latter to single-cell data. This method enables us to obtain expressions for the standard telegraph model parameters as a function of the size of extrinsic noise (the coefficient of variation of the parameter that varies the most between cells) and of the actual population averaged parameters. We show that (i) the estimated parameters are not generally equal to the population average of these parameters; (ii) whenever three moments of the telegraph model can be fitted to those of the data, i.e. none of the estimated parameters are negative, there are unique bias signatures depending on the source of extrinsic noise, i.e. which parameter is the most variable amongst cells. Variability in the synthesis rates leads to an overestimation of the three transcriptional parameters (synthesis rate and the two switching rates) and an underestimation of the burst size. Variability in the switching on rate has the exact opposite effect. Variability in the switching off rate leads to an overestimation of the synthesis rate and underestimation of the switching on rate; the switching off rate is mostly underestimated while the burst size can be over or underestimated depending on the speed of promoter switching; (iii) the size of bias is significantly diminished when extrinsic noise is in one of the switching rates and the promoter switches slowly between inactive and active states compared to the rate of mRNA degradation; (iv) independent of the magnitude of estimation bias, the mRNA count distributions of the standard telegraph model are in most cases practically indistinguishable from those of the telegraph model with extrinsic noise. This implies that given data from only one gene, standard model selection criteria (Akaike and Bayesian information criteria) would likely lead to the conclusion that the standard telegraph model is the model of choice even if the stochasticity in count data is not due to intrinsic sources; (v) extrinsic noise in the synthesis rate leads to a massive overestimation of the synthesis and switching off rates (but has little effect on the switching on rate) when the extrinsic noise size is close to a critical threshold. Beyond this critical size, it is not possible to match the first three moments of the telegraph model to data; (vi) for genes exhibiting transcriptional bursting, only the first two moments can be reliably matched. In this case extrinsic noise, independent of its source or distribution, will lead to an overestimation of the burst size and an underestimation of the burst frequency; (vii) while our analytical results assume moment-matching and an infinite sample size, we showed that similar numerical results are obtained by maximising the likelihood of the telegraph model using data with realistic sample sizes, a commonly used method.

We note that result (i) is not an obvious one. In fact given a parameter that varies about some mean from one cell to another, one would expect this noise to average out when performing inference using the standard telegraph model. This reasoning is implicit in the wide use of this model in smFISH and scRNA-seq studies. But our study shows that this averaging does not occur; rather extrinsic noise leads to a systematic estimation bias which artificially inflates some parameter values while having the opposite effect on others. A particular instance of these deviations was reported in a recent experimental study [13]. The telegraph model parameters were determined for the GAL10 gene for cells in the G1 phase, in the G2 phase and for cells in both phases together (merged data). It was found that the synthesis rate, the switching on rate and the switching off rates estimated using the merged data were larger than the true population averaged values, i.e. the weighted average of the parameters in the G1 and G2 phases. The opposite was true for the burst size. The sign of the bias is exactly the same as in one of our theoretical results (see result (ii) above). Of course one can argue that if the size of the bias is small then there would not be much cause for worry. But by result (v) we have that for common sources of extrinsic noise which affect the synthesis rate (such as the coupling of this rate and cell size to achieve concentration homeostasis [14]), the standard telegraph model estimates of the synthesis and switching off rates go to infinity as the size of extrinsic noise (the variability of the synthesis rates between cells) approaches a critical value. This could explain why a recent study [6] that estimated transcriptome-wide parameters using the standard telegraph model found a significant number of mammalian genes for which the synthesis and the switching off rates are a hundred to a thousand times larger than the switching on rate – examples of such genes are Sdpr, Sf3b1, Hspe1, Cyb5r1, Phlda3 and Prrx1. In fact our reanalysis of the scRNA-seq data for ∼2000 highly bursty mouse fibroblast genes in this study showed that taking into account extrinsic noise leads to burst frequency and size distributions that are remarkably different to those initially found using the standard telegraph model. While the count distributions for each gene are well fit by both the standard telegraph model and that with extrinsic noise (as found in result (iv)) however we showed that the two models can be told apart using the covariance of mRNA counts between different genes.

The main limitations of our study are: (i) the assumption that the mRNA degradation rate does not vary between cells. Relaxing this assumption makes it difficult to understand the effects of extrinsic noise on parameter estimation because it implies cell-to-cell variability in all three normalised transcriptional parameters of the telegraph model. Some studies have found that degradation rate does vary between cells [31, 63]. Other studies have found no relationship between cell size and degradation rate and thus posited that it is not a major source of extrinsic noise [14, 15]; (ii) the assumption that the data is single-allele. This data is nowadays readily available. However if one is interested in understanding the accuracy of inference from non-allele-specific data then a similar moment-based analysis as we did for one allele can be carried out provided one assumes independent expression from each allele [7, 13]; (iii) the assumption that each cell has a set of time-independent parameters. While generally it is to be expected that parameters vary with time inside a cell, nevertheless if the timescale of variation is slow enough then a cell will alternate between a number of steady states and a measurement will capture a cell in one of these – our theory is valid provided this is true. Moments of the standard model modified to account for dynamic extrinsic noise can be derived in specific cases and under some assumptions [64–66]. However because of the complexity involved, it will generally be difficult to extract relationships between the true population averaged parameters and those inferred; (iv) the correction of burst frequency and size from scRNA-seq data to account for extrinsic noise assumes that the UMI counts for a gene accurately represents the true counts. Whilst this has been assumed previously [6] it cannot be the case because typically the capture efficiency of sequencing technologies is not very high [53]. The capture efficiency model postulated in [59, 61] suggests that variability in capture efficiency between cells can be modelled by a variable synthesis rate. If this is indeed the case then our theory’s corrections for the population averaged burst frequency for highly bursty genes in [6] are accurate. On the other hand, our theory’s corrections for the population averaged burst size would depend on both cell-to-cell variation in the synthesis rate and in the capture efficiency and hence would not purely reflect the natural variation in parameters. Thus generally it is possible that the burst frequency is more accurately inferred than the burst size. It is also likely that explicit modelling of technical noise arising due to RNA-sequencing protocols is needed to understand how to properly correct the count data from scRNA-seq studies [67, 68].

The overall implication of our study is to avoid parameter estimation from marginal steady-state distributions of the mRNA counts using the standard telegraph model because the results are meaningless if there is considerable parameter variability between cells (the typical case). This type of estimation is only feasible if the major sources of extrinsic noise can be identified and the data is appropriately corrected for them prior to inference or else if the parameters are corrected *a posteriori*.

## AUTHOR CONTRIBUTIONS

R. G. designed research, performed research, contributed analytic tools, analyzed data, and wrote the paper. P-M. E. performed research and contributed analytic tools.

## ACKNOWLEDGMENTS

We thank Augustinas Sukys for useful feedback and help with the figures. This work was supported by a Leverhulme Trust research award (RPG-2020-327).

## Appendix A: Moments of the telegraph model with extrinsic noise

Let *x*^(*k*)^ = *x*(*x* + 1) (*x* + *k* 1) (resp. *x*_(*n*)_ = *x*(*x* − 1)⋯(*x* − (*k* − 1))) be the rising (resp. falling) factorial, and 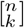 the absolute value of the corresponding Stirling number of the first kind. These numbers are related by the following identity:

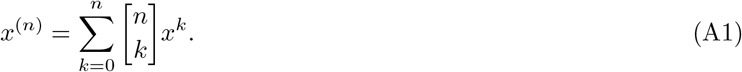

### 1. Derivation of the factorial moments of the telegraph model

The steady-state distribution of mRNA counts P(*m*) of the telegraph model (Eq. (1)) is exactly equal to the Beta-Poisson distribution, i.e. the distribution of a Poisson random variable with parameter *α*·*r*, where r is a random variable that is distributed according to the Beta(*β, θ*) distribution [35, 69]. Since the *n*th factorial moments of a Poisson distribution is the *n*th power of its parameter, it immediately follows that to obtain the *m*th factorial moment of the mRNA counts *m* of the telegraph model, we only have to calculate the following integral:

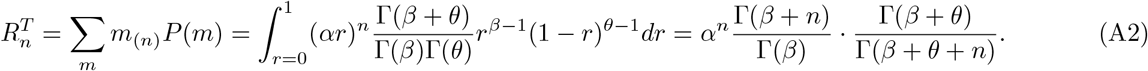

### 2. Derivation of the factorial moments of the telegraph model with gamma-distributed noise

To distinguish the models when there is extrinsic noise, we will consider the setup shown in Fig. 2(a)-(c) left panels where now we call the synthesis rate ρ, the switching on rate k_on_ and the switching off rate k_off_. We will consider each of these parameters to be distributed according to a gamma distribution with shape parameter a and scale parameter b.

#### a. A few useful formulas

Using a property of the gamma function, one can write the rising factorial as a quotient of two gamma functions:

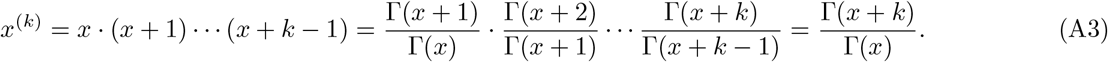

For *n* ≥ 1 the following partial fraction decomposition holds:

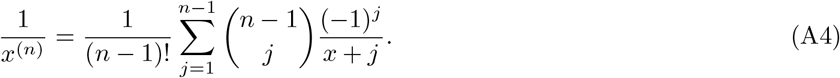

This can be derived by setting 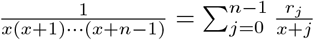 where

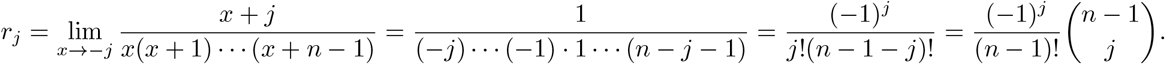

Kummer’s function of the second kind *U*(*a, c, z*) can be expressed as [70]:

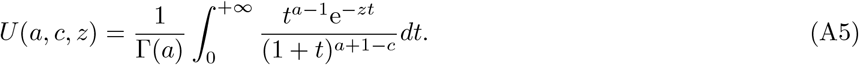

For *a, b, α* > 0 the following equalities hold:

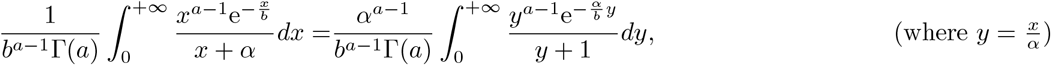

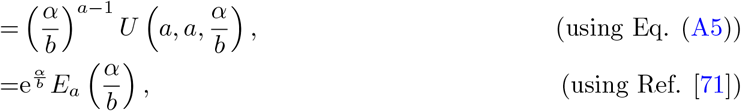

where 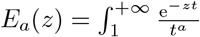 dt is the exponential integral function. Hence

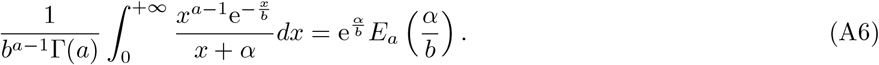

#### b. Gamma noise on the synthesis rate ρ

To get the factorial moments in this case, we integrate Eq. (A2) (after relabelling the parameters as described in the beginning of this section) over a gamma distribution in the synthesis rate:

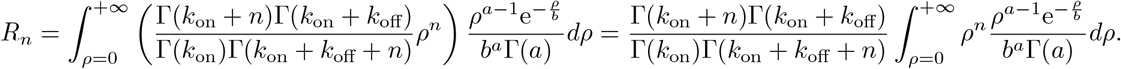

Using the normalisation of the gamma distribution, we immediately have 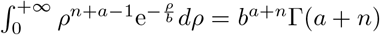, which gives the following result:

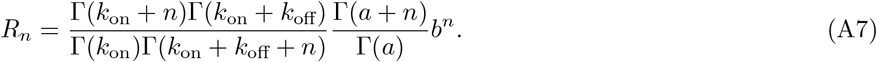

This result has been previously derived in [34].

#### c. Gamma noise on the switching on rate k_on_

To get the factorial moments in this case, we integrate Eq. (A2) over a gamma distribution in the switching on rate. The integral can be simplified by using Eq. (A3) to write the two quotients of gamma functions in terms of rising factorials:

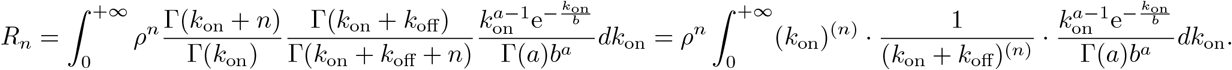

Next using Eqs. (A1) and (A4) we get rid of the rising factorials

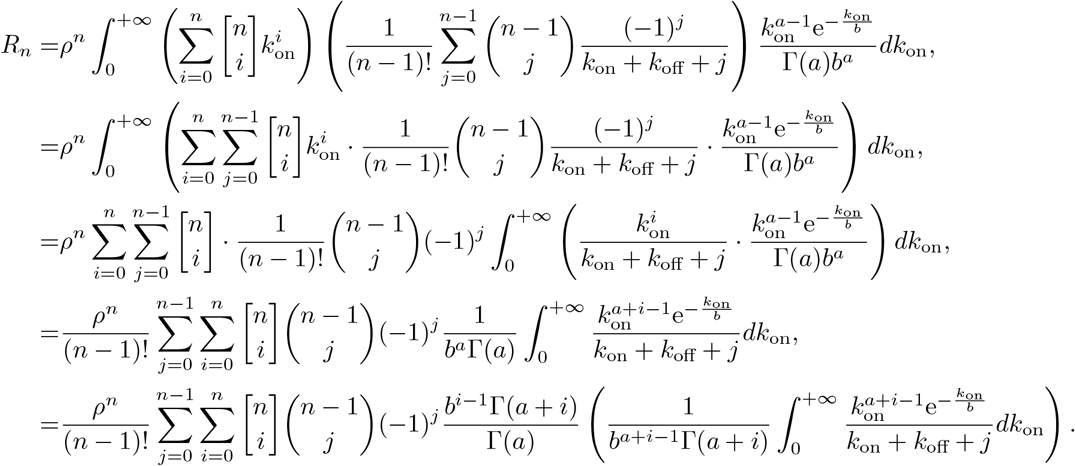

Finally using Eq. (A6) we obtain

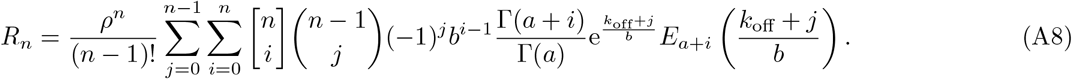

#### d. Gamma noise on the switching off rate k_off_

To get the factorial moments in this case, we integrate Eq. (A2) over a gamma distribution in the switching off rate. The integral can be simplified by replacing the quotient 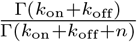 by a rising factorial (using Eq. (A3)):

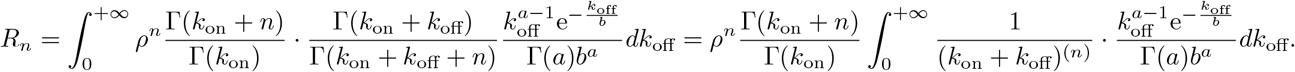

Simplifying further using Eq. (A4) (to get rid of the rising factorial), leads to

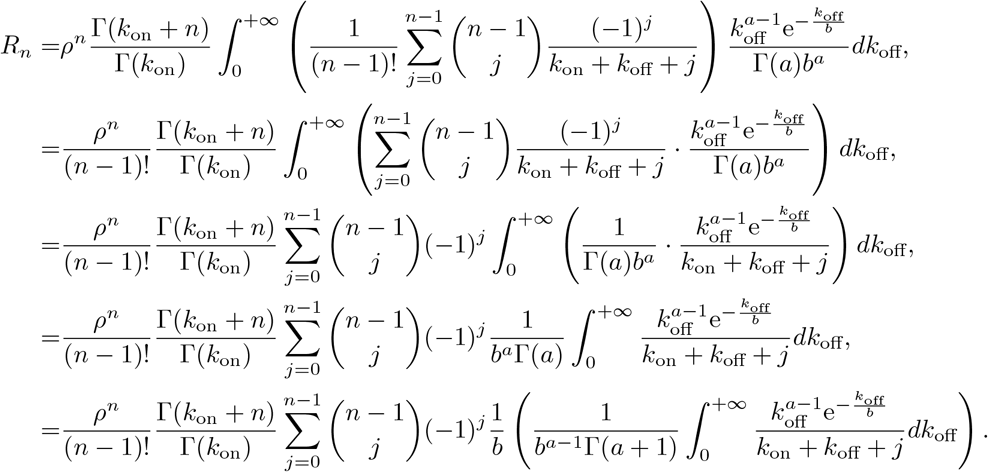

Finally by using Eq. (A6) we find

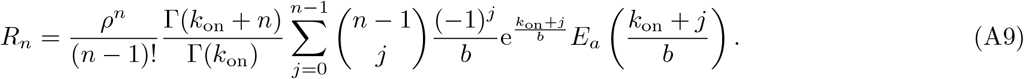

## Appendix B: Estimation bias in the limit of small extrinsic noise

In this section, we study estimation bias when the experimental data can be described by the telegraph model with small extrinsic noise, i.e. the limit of small cell-to-cell variability in parameter values. Our solution strategy is as follows. We will compute the factorial moments of mRNA counts for the telegraph model with small extrinsic noise by means of Taylor’s approximation for moments [72]. For simplicity, we will consider only one parameter to vary amongst cells. Let this random parameter 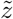 have mean z and a small standard deviation *σ*_*z*_. We do not make further assumptions on the type of distribution of this parameter. By the aforementioned approximation for some function *f* of the random parameter, we have

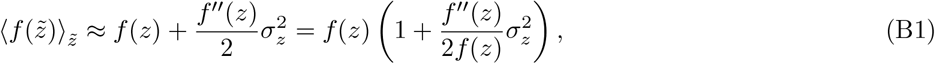

where the angled brackets is understood to mean averaging over the distribution of the random parameter. Then we will equate the factorial moments of mRNA counts for the telegraph model with small extrinsic noise with the factorial moments of the standard telegraph (mimicking inference using this model) and solve them for the telegraph model parameters {*α, β, θ*} using Eq. (2). Finally by taking the derivative of these parameters with respect to 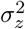 when *σ*_*z*_ = 0, we will determine the sign of estimation bias.

### 1. General theory

For the sake of generality, the beginning of the derivation will use a tilde over all three parameters. Later on we will consider only one parameter to be noisy, and the other two to be equal to their mean.

The first three factorial moments of mRNA counts as predicted by the standard telegraph model are given by (Appendix A):

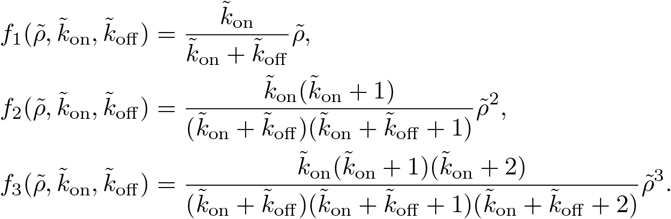

By applying Taylor’s approximation Eq. (B1) on the functions *f*_*i*_ we obtain the extrinsic noise averaged factorial moments:

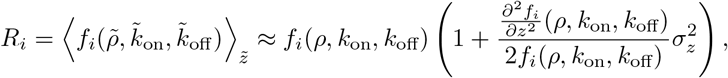

where *z* can be *ρ, k*_on_ or *k*_off_. By means of the notation 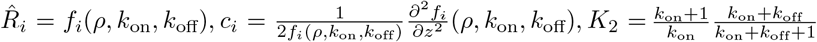 and 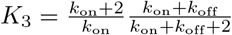, these expressions can be compactly written as

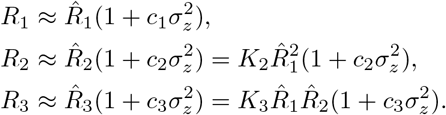

Substituting these in Eq. (2) we find the telegraph model estimates of the three transcriptional parameters:

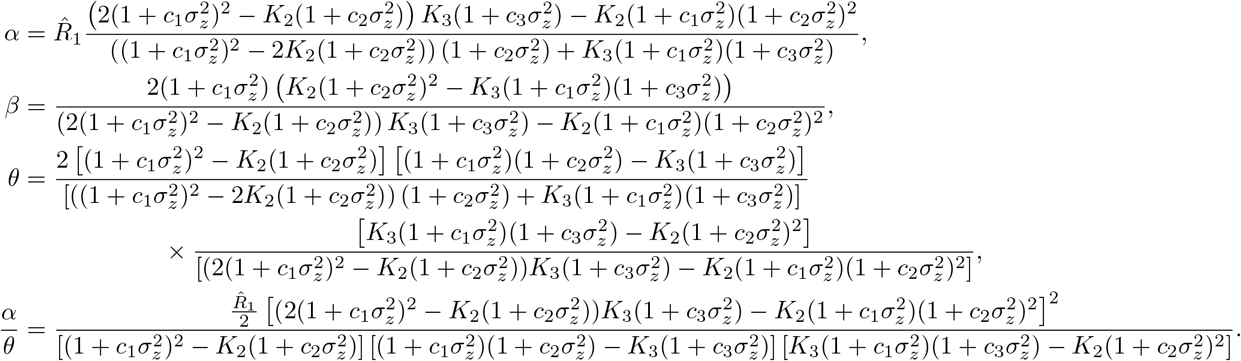

It is easy to show that if 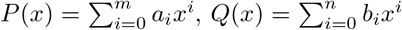 and 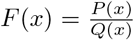, then 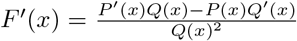 and 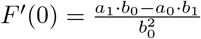.

Using this result, the derivatives with respect to 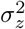, evaluated at 0, are given by

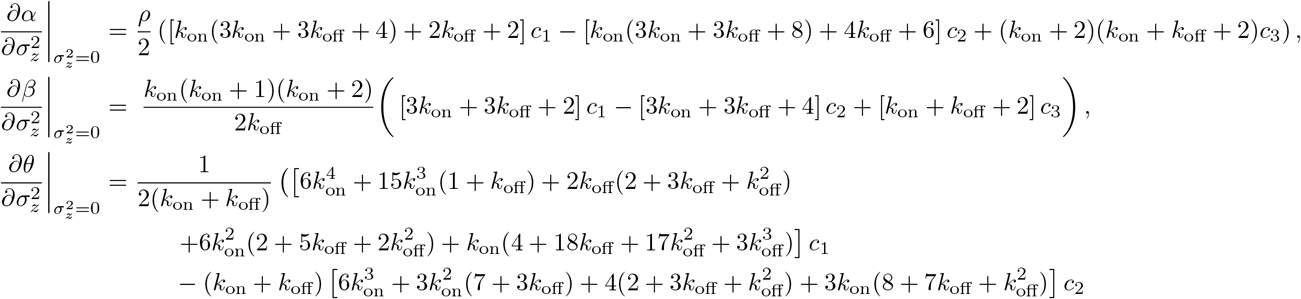

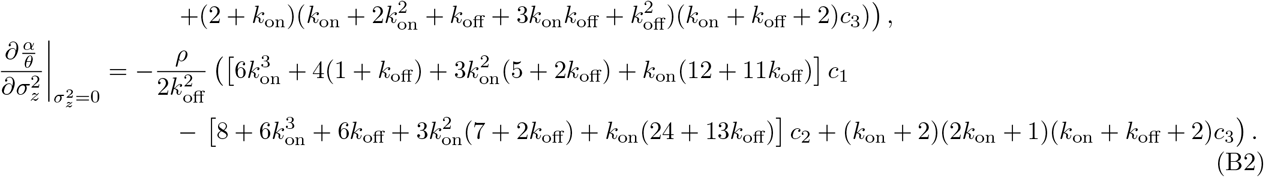

### 2. Variable synthesis rate: 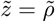

By the definition of the formulae for c_*i*_, we have:

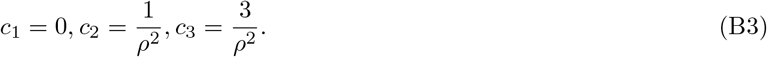

Substituting in Eq. (B2) we obtain:

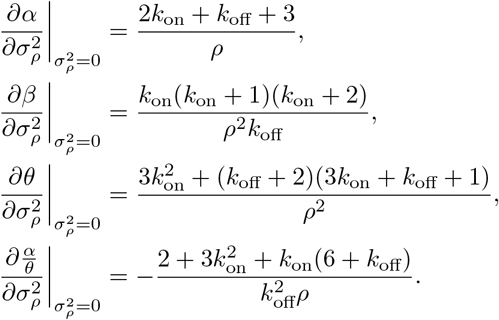

Hence it follows that in the presence of small extrinsic noise in the synthesis rate, use of the telegraph model for inference will lead to an overestimation of the synthesis, switching on and switching off rates and an underestimation of the burst size. This agrees with Fig. 2(a).

### 3. Variable switching on rate: 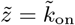

By the definition of the formulae for c_*i*_, we have:

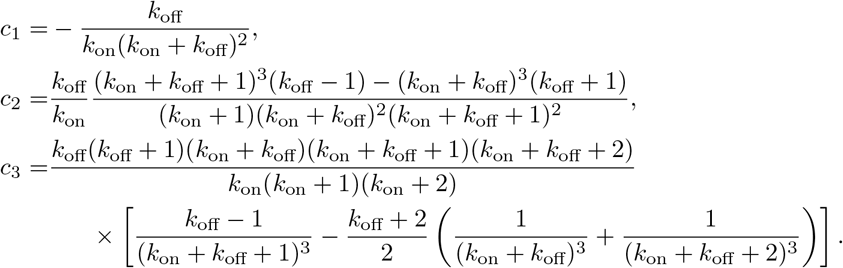

Substituting in Eq. (B2) we obtain:

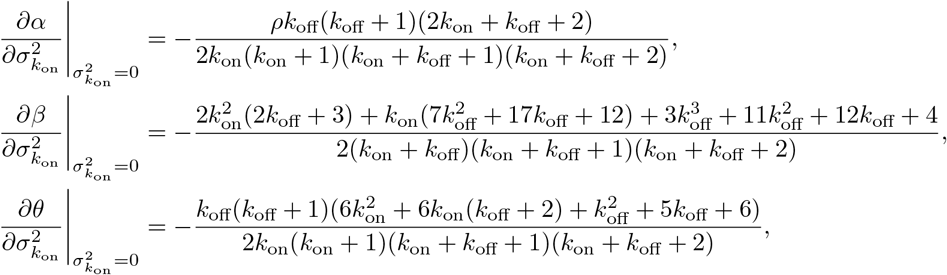

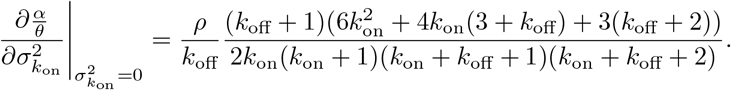

Hence it follows that in the presence of small extrinsic noise in the switching on rate, use of the telegraph model for inference will lead to an underestimation of the synthesis, switching on and switching off rates and an overestimation of the burst size. This agrees with Fig. 2(b).

### 4. Variable switching off rate: 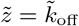

By the definition of the formulae for c_*i*_, we have:

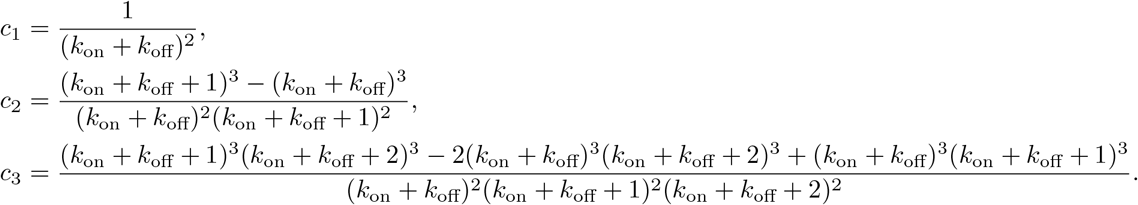

Substituting in Eq. (B2) we obtain:

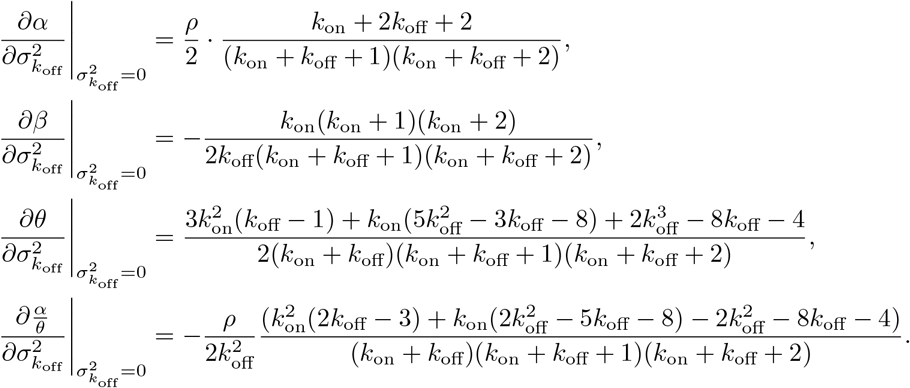

The first and second expressions are respectively positive and negative, implying that the telegraph model overestimates the synthesis rate and underestimates the switching on rate. However the third and the fourth expressions can be positive or negative, depending on the parameter values, and hence both over and underestimation are possible for the switching off rate and the burst size. This agrees with Fig. 2(c). Comparing the box-whisker plots in the middle and right columns, it is clear that the sign of the estimation bias for the burst size changes if promoter switching is constrained to be slow. This is also the case for the switching off rate but cannot be appreciated from the plots (the maximum value of *θ*/ ⟨*k*_off_⟩ is slightly greater than 1 but the upper fence of the box-whisker plot only shows the 90th percentile).

The results of the small-noise theory are summarised in Table II.

**TABLE II.** Summary of the small extrinsic noise theory showing the type of bias in the telegraph model parameters (*α, β, θ, α/θ*) when these are inferred from data generated by a telegraph model with small extrinsic noise in one of the parameters (*ρ, k*_on_, *k*_off_).

| | Noise on $\rho$ | Noise on $k_{\text{on}}$ | Noise on $k_{\text{off}}$ |
| --- | --- | --- | --- |
| $\alpha$ | overestimation | underestimation | overestimation |
| $\beta$ | overestimation | underestimation | underestimation |
| $\theta$ | overestimation | underestimation | indeterminate |
| $\frac{\alpha}{\theta}$ | underestimation | overestimation | indeterminate |

## Appendix C: Parameter details for Figure 3

The telegraph model distributions of mRNA counts (solid blue lines in the figure) are evaluated using the analytical steady-state solution [1] for the parameters estimated from the three-moment matching procedure.

For (a), the parameters of the telegraph model with extrinsic noise are ⟨*ρ*⟩ = 3.92872, *k*_on_ = 2.74898, *k*_off_ = 5.20897, 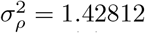 and the inferred parameters of the telegraph model are *α* = 26.2939, *β* = 2.87887 and *θ* = 52.8984.

For (b), the parameters of the telegraph model with extrinsic noise are *ρ* = 10.4132, ⟨*k*_on_⟩ = 179.703, *k*_off_ = 86.4225, 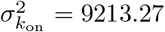 and the inferred parameters of the telegraph model are *α* = 8.70773, *β* = 4.88633 and *θ* = 1.56185.

For (c), the parameters of the telegraph model with extrinsic noise are *ρ* = 30.2479, *k*_on_ = 20.8791, ⟨*k*_off_⟩ = 49.9974, 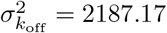 and the inferred parameters of the telegraph model are *α* = 50.5625, *β* = 2.12659 and *θ* = 6.47778.

## Appendix D: Estimation bias when the synthesis rate is lognormal distributed

Consider a population of cells where the synthesis rate *ρ* varies from cell to cell. Furthermore we assume that this variation is described by a lognormal distribution with parameters (*µ*, s). The mean of this distribution is 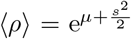, and the second moment is 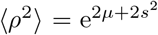. The coefficient of variation squared of the synthesis rate is equal to 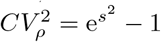. The factorials moments of mRNA counts can be straightforwardly computed [34]

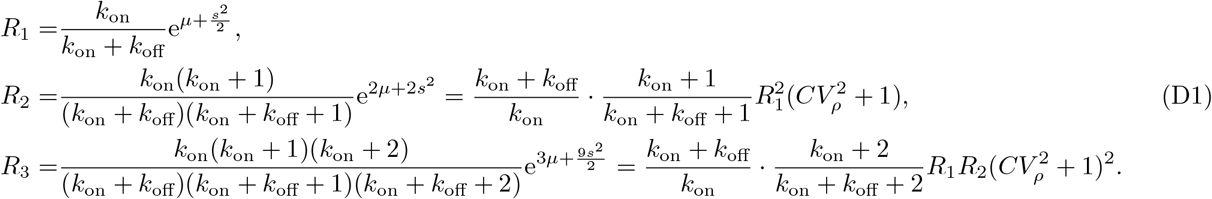

Substituting in Eq. (2), we obtain the parameters inferred by fitting a telegraph model:

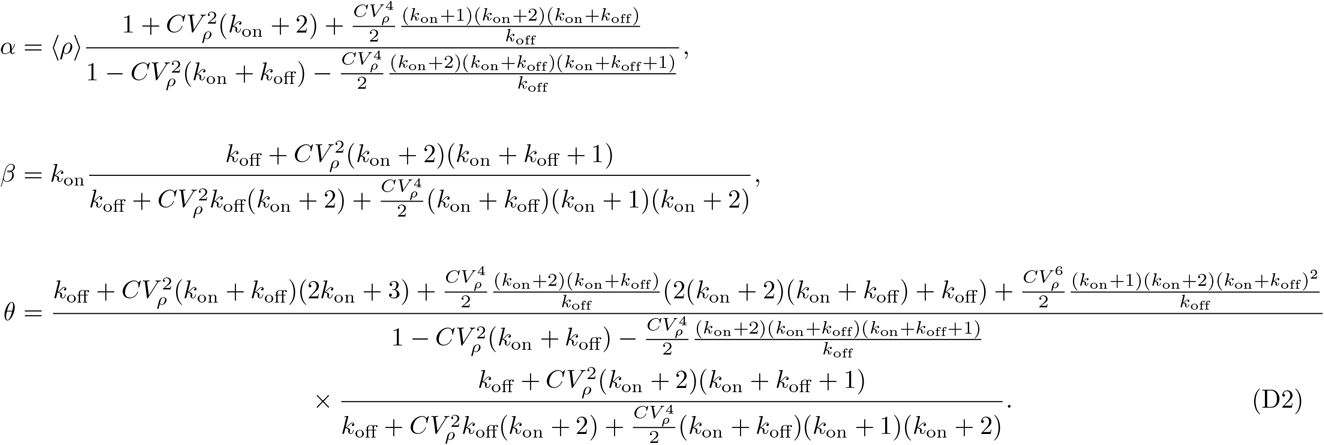

It is immediately evident that β is always positive, and that the sign of the other two parameters only depends on the sign of their denominator. The discriminant of the denominator is

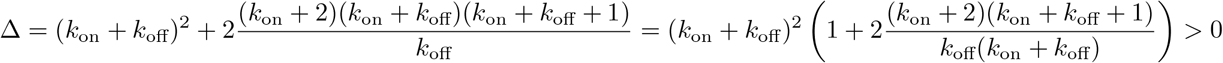

and the two solutions are given by

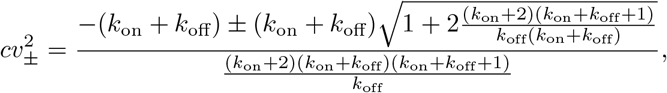

where 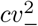 is negative and 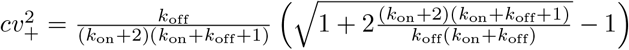 is positive. Hence the parameters α and θ are positive provided the coefficient of variation squared of the synthesis rate 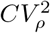 is less than 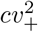.

If we compare this critical noise threshold with that for gamma-distributed noise (Section III) we find that it is always smaller

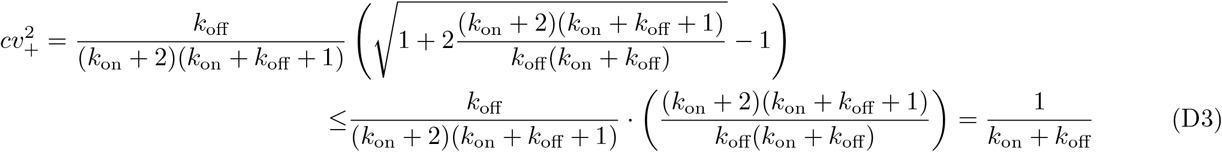

A summary of the properties of the estimated rates is as follows:

- *α* is positive only when the extrinsic noise size is below the threshold value 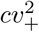. In this case it is an increasing function of 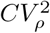 and always larger than ⟨*ρ*⟩. It diverges as 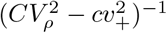 near the critical noise threshold.
- *β* is always positive, independent of the size of the extrinsic noise. It increases monotonically with the extrinsic noise size on 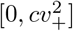; in this region, it is larger than *k*_on_ and reaches its maximum when 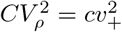.
- *θ* is positive only when the extrinsic noise size is below the threshold value 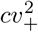. In this case it is an increasing function of 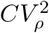 and always larger than *k*_off_. It diverges as 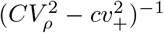 near the critical noise threshold.

## Appendix E: Maximum likelihood estimation and the generation of simulated data

### 1. Maximum Likelihood estimation using the standard telegraph model

Given the steady-state distribution of mRNA counts predicted by the standard telegraph model

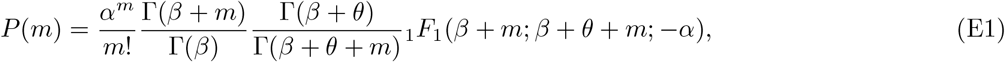

and the data vector 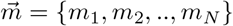 (where *m*_*i*_ is the mRNA counts in cell *i*), the procedure consists of finding the parameter set {*α, β, θ*} that maximises the log-likelihood function defined as

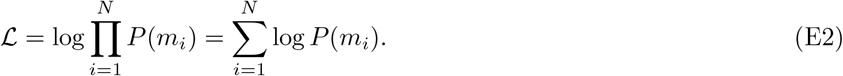

The optimisation was performed using Mathematica’s FindMaximum function. Good initial guesses for the optimisation procedure can sometimes speed up estimation [46]; for our numerical experiments, we did not find a large improvement in speed and the results were the same if we simply constrained the optimisation to search over positive parameter values.

### 2. Simulating mRNA count data

To mimic experimental mRNA count data, we need to sample from the steady-state distribution of the telegraph model with extrinsic noise. For this purpose we made use of the equivalence between the standard telegraph model’s solution and the Beta-Poisson compound distribution (discussed in Appendix A). For example if we want to generate data for the telegraph model with gamma distributed synthesis rate (Fig. 2(a)), the mRNA count for each cell was obtained by drawing a Poisson distributed random number Poisson(*r*_1_*r*_2_) where r_1_ is a random number Beta(*k*_on_, *k*_off_) and r_2_ is a random number Gamma 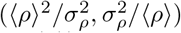. Similarly to generate data for the telegraph model with gamma distributed switching on rate (Fig. 2(b)), the mRNA count for each cell can be obtained by drawing a Poisson distributed random number Poisson(*ρr*_1_) where *r*_1_ is a random number Beta(*r*_2_, *k*_off_) and *r*_2_ is a random number Gamma 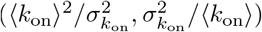. Note that ⟨*i*⟩ and 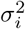 denote the mean and variance of parameter *i*, respectively.

## Appendix F: Covariance matrix analysis of scRNA-seq data

### 1. Extrinsic noise on the synthesis rate due to coupling of transcription to cell volume

Consider a a population of cells characterized by a distribution *f*(*V*) of the cell volume. Furthermore consider the stochastic expression of two genes *G*_1_ and *G*_2_ whose expression dynamics in a cell of volume *V* is described by the following modified telegraph model:

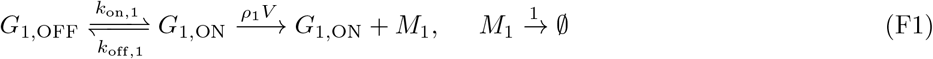

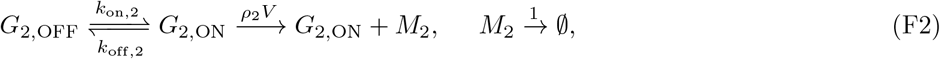

where *M*_*i*_ is mRNA from gene *G*_*i*_. Note that the rates for each gene are normalised by the mRNA degradation rate for that gene (hence the latter is set to 1).

We now derive the expression of the covariance Cov(*m*_1_, *m*_2_) between the mRNA counts *m*_1_ and *m*_2_ of the two genes. Given a cell with volume *V*, we have the following moments:

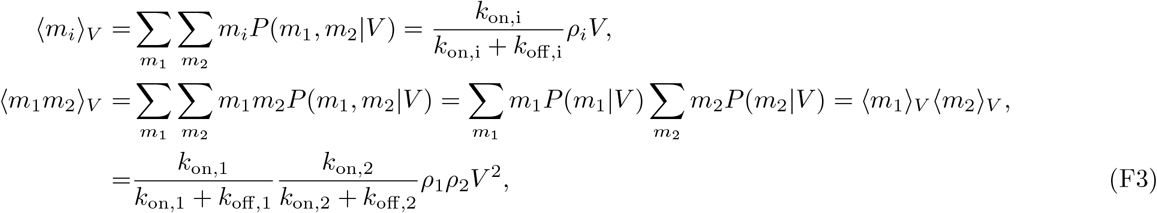

where we have used the independence of the two genes in the last line. Hence it follows that by taking the average over the extrinsic noise, i.e. integrating over the cell size distribution, we obtain:

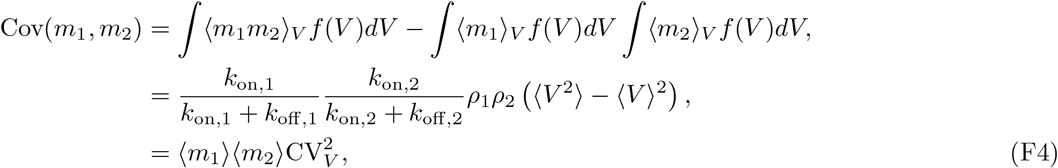

where ⟨*m*_*i*_⟩ is the integral of ⟨*m*_*i*_⟩_*V*_ over the volume distribution *f*(*V*) and 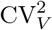 is the coefficient of variation of the volume distribution. It is interesting that extrinsic noise, i.e. variability in the synthesis rate due to variability in the cell volume, causes the covariance to be always positive. Hence plotting the covariance of two genes versus the product of their mean expression should lead to a straight line with a slope equal to the size of extrinsic noise; in Fig. 6(b) we confirm this using the scRNA-dataset in [6].

### 2. Extrinsic noise on the burst size or frequency for bursty genes

We consider some source of extrinsic noise (cell-specific) which acts linearly on a bursty gene either via modulation of its burst size or burst frequency. We will denote this global factor by *V* (does not necessarily mean volume).

For noise in the burst size, the results following directly from those above. The burst size is the synthesis rate divided by the switching off rate hence noise in the synthesis rate will lead to noise in the burst size. Thus Eq. (F4) immediately follows.

For noise in the burst frequency and transcriptional bursting, we consider the reaction scheme (F2) with the changes: *ρ*_*i*_*V* → *ρ*_*i*_, *k*_on,i_ →*k*_on,i_*V*. Then taking the limit of transcriptional bursting, i.e. *k*_on,i_*V*_*max*_ ≪*k*_off,i_, one finds that ⟨*m*_*i*_⟩ _*V*_ ∝ *V* and ⟨*m*_1_*m*_2_⟩ _*V*_ ∝*V*^2^ which implies that Eq. (F4) holds. Note that *V*_*max*_ is the maximum value of the extrinsic parameter *V*.

## Appendix G: Normalisation of scRNA-seq data can introduce estimation bias

In [61], for a given gene, a telegraph model with extrinsic noise in the synthesis rate is proposed to describe its stochastic expression and to take into account the capture-efficiency of scRNA-seq technology. Expression from cell *j* is assumed to be described by reaction scheme in Fig. 2(a) left with the change *ρ* →*ργ*_*j*_ where *γ*_*j*_ is a cell-specific factor that is the product of the capture efficiency and the volume for cell j. A recipe is then described for computing *γ*_*j*_ for each cell from the raw scRNA-seq data. Given these cell-specific factors and mRNA count data from each cell their goal is to infer the switching rates *k*_on_ and *k*_off_, and the parameter *ρ*.

One of the heuristic methods proposed involves a modification of the moment-matching procedure we described in Section II. It is proposed that one uses Eq. (2) with the difference that the *i*th factorial moment *R*_*i*_ is computed not for the observed counts data but rather for the counts data normalised by the cell-specific factor, i.e. m_*i*_ (observed counts in cell *i*) is replaced by *x*_*i*_ = *m*_*i*_/γ_*i*_. Since we can sample from the steady-state distribution of the telegraph model using the Beta-Poisson distribution, it follows that for cell *i* the normalised counts are given by

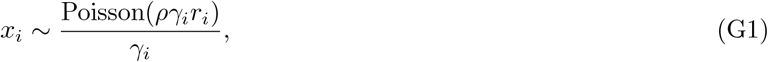

where *r*_*i*_ is a random variable distributed according to Beta(*k*_on_, *k*_off_). The heuristic method then is equivalent to assuming that the moments of *x*_*i*_ are precisely the same as the moments of the random variable

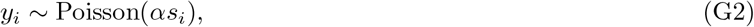

where *s*_*i*_ is a random variable distributed according to Beta(*β, θ*). Perfect inference by the heuristic method would imply *α* = *ρ, β* = *k*_on_ and *θ* = *k*_off_. By inspection of Eqs. (G1) and (G2), this only occurs if the following statement is true:

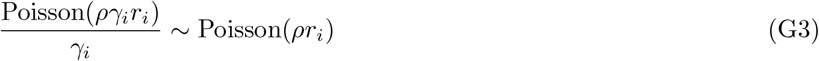

It is clear that this cannot be generally true for any distribution of the cell-specific factor *γ*_*i*_. In fact it is not even true if *γ*_*i*_ is a constant independent of i. However the equivalence is achieved asymptotically in the limit of large *ρ*γ_*i*_*r*_*i*_ and large *ρr*_*i*_ since then the Poisson random variables can be replaced by their mean, i.e. when the mean expression level of a gene is high enough, the heuristic moment-matching method of [61] will give reliable results. In Table III we numerically show that the accuracy increases with the mean expression level thus validating our theoretical results.

**TABLE III.**
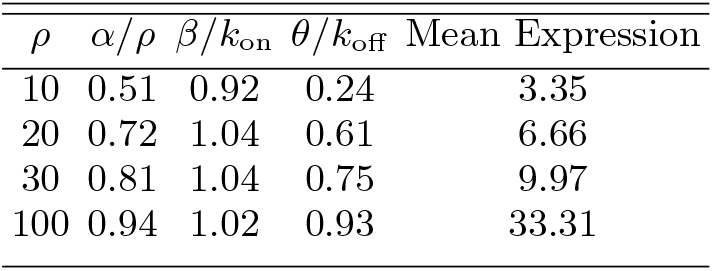
Numerical investigation of the accuracy of the parameter inference method in [61] based on moment-matching of the telegraph model to the moments of normalised mRNA counts. We simulated data for 10^5^ cells by sampling from the Beta-Poisson distribution with synthesis rate *ργ*_*i*_, switching on rate *k*_on_ = 1 and switching off rate *k*_off_ = 2, where *γ*_*i*_ is a gamma-distributed random variable with mean 20 and coefficient of variation squared = 1. The count data from cell *i* is then divided by *γ*_*i*_ leading to normalised counts. Substituting the first three factorial moments of the latter in Eq. (2) we obtain parameter estimates from the telegraph model (*α, β, θ*). Perfect parameter inference would mean *α* = *ρ, β* = *k*_on_, *θ* = *k*_off_. We vary the mean expression by increasing *ρ* whilst keeping all other parameters constant. In agreement with our theory (summarised in Eq. (G3)), the accuracy of the heuristic inference method increases with the mean expression.

We note that the errors in this method can be overcome by using a different moment-matching method. Under the assumption that scRNA-seq data is accurately described by a telegraph model with extrinsic noise on the synthesis rate (as assumed in the beginning of this section and in [61]) then exact parameter inference can be implemented by equating the first three moments of this model (derived for any distribution of extrinsic noise in the synthesis rate in [34]) with the first three raw moments of count data and then solving the three resultant equations simultaneously for ρ, k_on_, k_off_. If gene expression is bursty then this procedure is equivalent to the method described in Section IV.C with the burst size for gene i in cell j proportional to the cell-specific factor γ_*j*_. Of course the accuracy of this method relies on whether scRNA-seq data truly suffers only from noise on the synthesis rate (although generally unlikely this may be true for the data in [6]) and also on the estimation of the cell-specific factors γ_*j*_.

